# Diversity in Chemical Subunits and Linkages: A Key Molecular Determinant of Microbial Richness, Microbiota Interactions, and Substrate Utilization

**DOI:** 10.1101/2024.08.12.607625

**Authors:** Hugh C. McCullough, Hyun-Seob Song, Jennifer M. Auchtung

**Affiliations:** Nebraska Foods for Health Center, University of Nebraska-Lincoln, Lincoln, NE USA; Department of Food Science and Technology, University of Nebraska-Lincoln, Lincoln, NE USA; Department of Biological Systems Engineering, University of Nebraska-Lincoln, Lincoln, NE USA

## Abstract

Dietary fibers play a significant role in shaping the composition and function of microbial communities in the human colon. Our understanding of the specific chemical traits of dietary fibers that influence microbial diversity, interactions, and function remains limited. Towards filling this knowledge gap, we developed a novel measure, termed Chemical Subunits and Linkages (CheSL) Shannon diversity, to characterize the effects of carbohydrate complexity on human fecal bacteria cultured *in vitro* under controlled, continuous flow conditions using media that systematically varied in carbohydrate composition. Our analysis revealed that CheSL Shannon diversity demonstrated a strong Pearson correlation with microbial richness across multiple fecal samples and study designs. Additionally, we observed that microbial communities in media with higher CheSL Shannon diversity scores exhibited greater peptide utilization, and more connected, reproducible structures in computationally inferred microbial interaction networks. Together, these findings demonstrate that CheSL Shannon diversity can be a useful tool to quantify the effects of carbohydrate complexity on microbial diversity, metabolic potential, and interactions. Furthermore, our work highlights how robust and stable community data can be generated by engineering media composition and structure. These studies provide a valuable framework for future research on microbial community interactions and their potential impacts on host health.

**IMPORTANCE:** For the human adult gut microbiota, higher microbial diversity strongly correlates with positive health outcomes. This correlation is likely due to increased community resilience that results from functional redundancy that can occur within diverse communities. While previous studies have shown that dietary fibers influence microbiota composition and function, we lack a complete mechanistic understanding of how differences in the composition of fibers are likely to functionally impact microbiota diversity. To address this need, we developed Chemical Subunits and Linkages (CheSL) Shannon diversity, a novel measure that describes carbohydrate complexity. Using this measure, we were able to correlate changes in carbohydrate complexity with alterations in microbial diversity and interspecies interactions. Overall, these analyses provide new perspectives on dietary optimization strategies to improve human health.

## INTRODUCTION

Members of the gut microbiota are known to contribute to different aspects of human health, including metabolizing otherwise inaccessible dietary substrates, educating the immune system, and providing protection from pathogen infection (1–4). Ongoing research efforts focus on developing treatments and interventions to engineer the gut microbiota to improve host health (5–8); approaches have included addition of new dietary substrates (e.g., prebiotics and dietary fibers (9–11)), new microbes (e.g., probiotics and live microbial therapeutics (1, 12, 13)), or combinations of substrates and microbes (e.g., synbiotics (2, 14)). Previous studies have shown diets are a major driver of microbial composition (15). In mice, consumption of a westernized diet caused large changes in microbiota composition compared to consumption of a low fat, low sugar diet enriched in complex polysaccharides (16, 17). Similarly, long-term dietary patterns and microbiota composition are correlated in humans (18). Dietary fibers are of particular interest for gut microbiota modulation because they cannot be metabolized by human enzymes and gut microbiota metabolism of these substrates can produce products (e.g., butyrate) metabolized by the host (19–22).

The “nutrient-niche theory” proposed by Freter et al. (23, 24) can serve as a basic framework to understand the impacts of diets on microbiota composition. It states that the metabolic potential of microbial community members and the availability of nutrients dictate how niches are occupied (23, 24). Particularly, increasing the number of unique utilizable substrates expands the fundamental niche space potentially facilitating increased microbial richness. However, competition for limiting nutrients and environmental conditions that limit growth may restrict the ability of microbes to colonize (25). Recent work has begun to investigate how carbohydrate structural diversity impacts microbial community composition. Chung et al. demonstrated that more complex fibers and/or mixtures of fibers increased microbial community richness and diversity compared to simple fibers for communities of human fecal microbes, although there was some variation between the two fecal communities tested (26). Yao et al observed similar effects culturing human fecal microbiotas, with the more complex fiber sorghum arabinoxylan supporting higher microbial community richness and diversity than the simple fiber inulin or mixtures of the primary monosaccharides found in either fiber (27). However, a follow-up study observed that the subtle differences in carbohydrate structure could have significant impacts on microbial richness of fecal communities cultured under high dilution pressure (28). Using a defined community of ten fecal microbes, Loss et al demonstrated that higher carbohydrate complexity and low sugar concentrations led to higher community diversity by increasing the ratio of positive to negative interactions between strains (29).

In this study, we analyzed the response of human fecal microbial communities to changes in the diversity and concentration of carbohydrates through a combination of *in vitro* experiments and computational modeling using a previously described continuous flow model of the lumen of the distal colon (30). Our workflow was composed of three parts (Fig. 1): (1) data generation and analysis to identify key traits of carbohydrates determining microbial richness, (2) post-analysis of metabolic potential and activity, and (3) network inference to predict community structure - microbial interactions and central microbiota. Inspired by the nutrient-niche theory, we developed a molecular-level diversity metric of carbohydrates termed Chemical Subunit and Linkage (CheSL) Shannon diversity, which we used to quantify carbohydrate diversity for mixtures of carbohydrates based on their internal linkages and subunits. Communities cultured in media with high CheSL Shannon diversity demonstrated strong positive correlations with microbial richness, peptide utilization (measured and predicted through Stickland fermentation), and microbial interactions in computationally inferred networks. Through these combined data analyses and model predictions, we also determined that microbial communities assembled under conditions with higher CheSL Shannon diversity led to more reproducible interactions with conserved central taxa (*Bacteroides*, *Lachnospiraceae* and *Ruminococcaceae*). Finally, we were able to draw parallels between enrichment of specific microbes in the presence of carbohydrate mixtures that were consistent with predicted abundances of metabolic pathways and enrichment previously observed *in vivo*.

**Figure 1:**
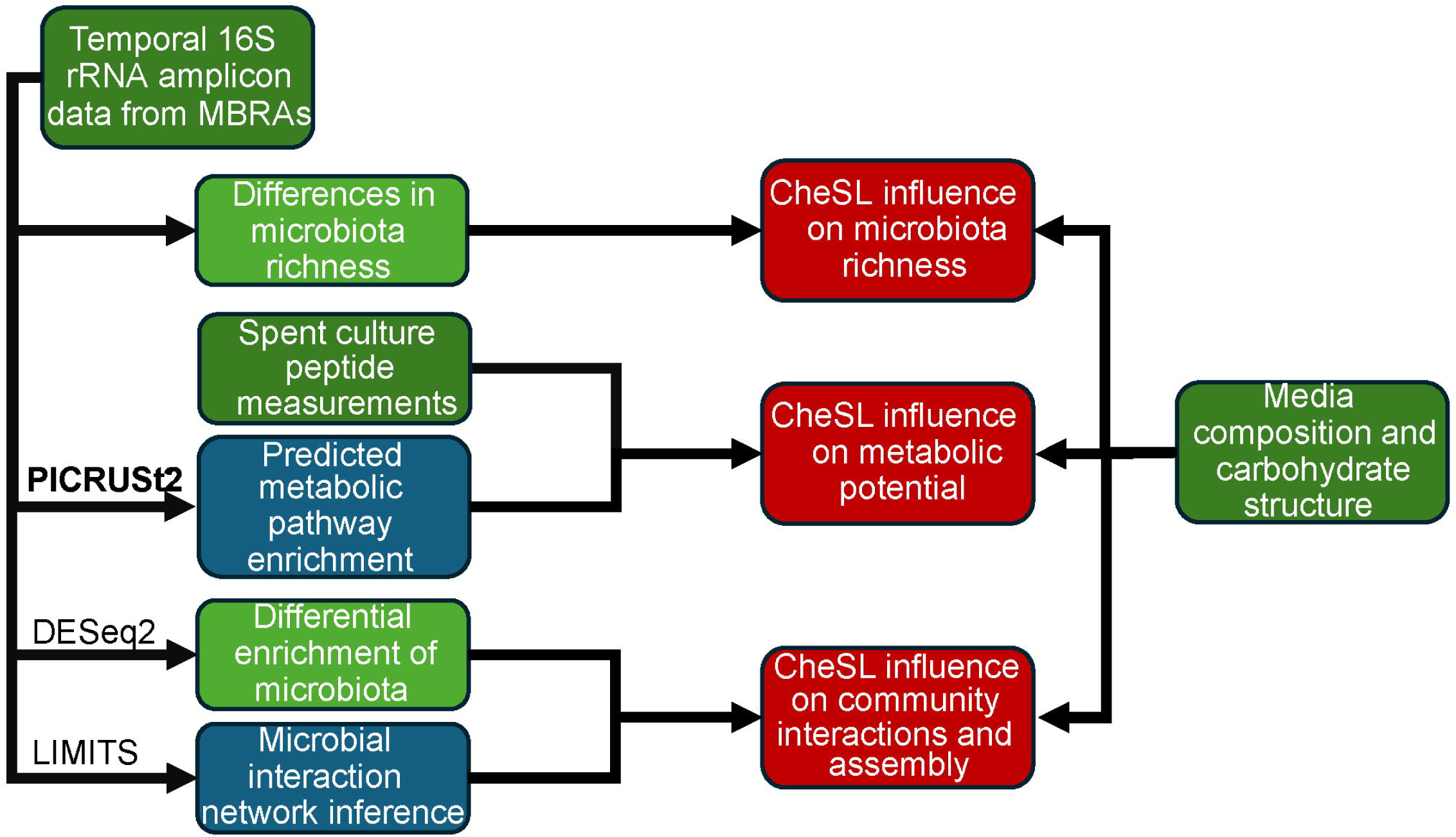
Graphical Abstract. A conceptual overview of this study, centered on applicability of CheSL metrics to different community culture traits. Empirical data is shown in dark green, post-analysis traits are shown in light green, inferred characteristics are show in blue, and the interpretations are shown in red.

## RESULTS

### Developing Metrics to Quantify Carbohydrate Complexity

To investigate the potential contribution of carbohydrate complexity to unique niches within a medium, we developed a new metric termed CheSL, which enables quantifying carbohydrate complexity based on the number of unique free monosaccharides and linkages present in each carbohydrate. Accounting for internal linkages in evaluating carbohydrate complexity is critical because polysaccharides may provide one or more unique niches depending on their complexity and the likelihood that enzymes required for degradation are encoded within a single microbe may vary based upon linkage complexity. For example, previous studies have demonstrated that while *Bifidobacterium longum* (31) and *Bacteroides thetaiotaomicron*, two common members of the gut microbiota, possess enzymes necessary for complete degradation of the simple fiber starch (32), neither *B. longum* nor *B. caccae* can fully metabolize arabinogalactan, a more complex polysaccharide composed of five different types of glycosidic bonds that connect primarily arabinose and galactose monomers (Table 1) (33).

**Table 1:**
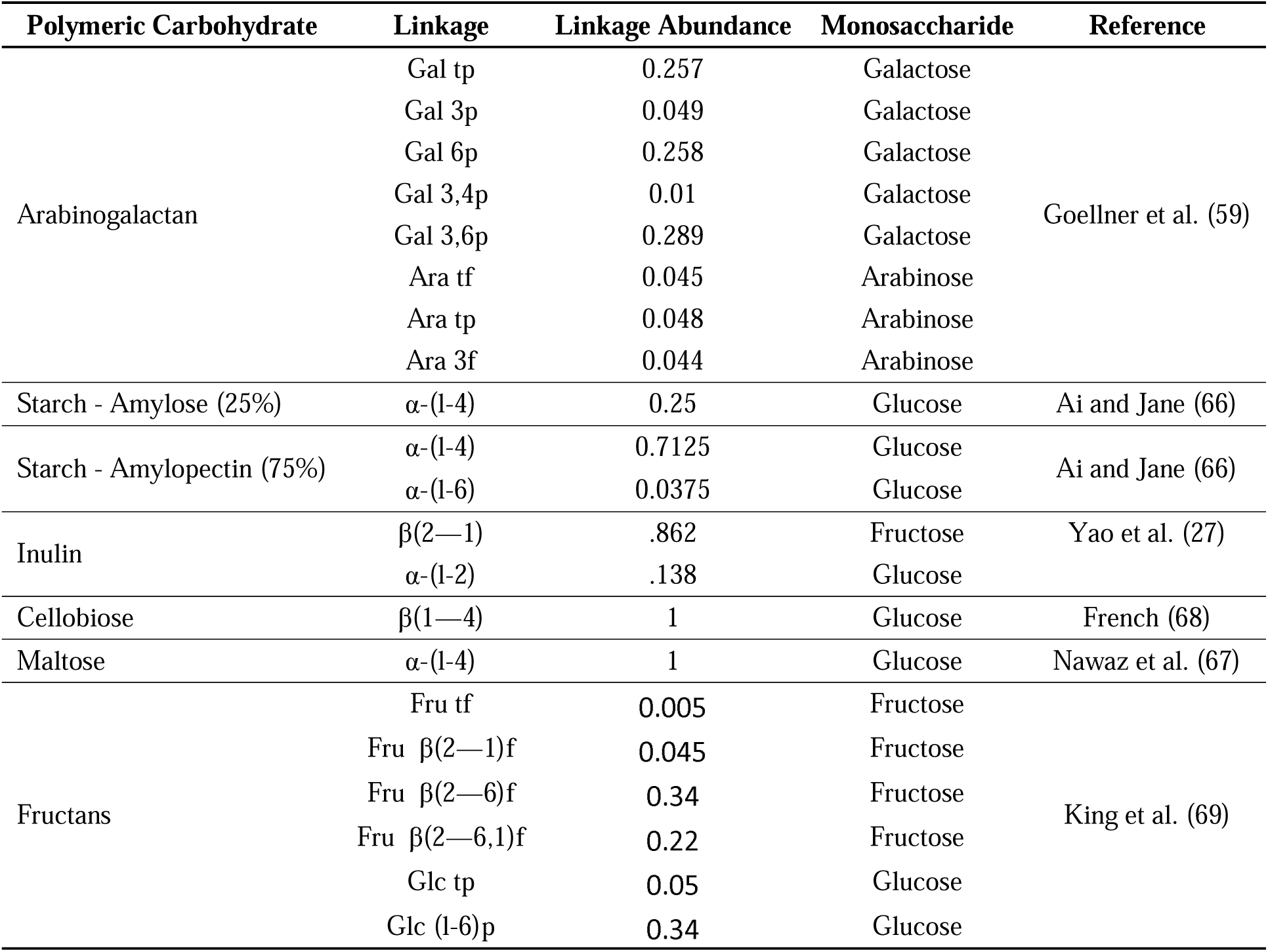
Linkages of Disaccharides and Polysaccharides tested. Linkages from the different polysaccharides used in this study, with estimated linkage abundances from literature, and in the case of starch, fractions of amylose and amylopectin determined as described in methods. For linkages, t indicates terminal, 3,4,6, refers to which carbon the bond is on, f indicates it is in the furanose form and p, the pyranose form.

| Polymeric Carbohydrate | Linkage | Linkage Abundance | Monosaccharide | Reference |
| --- | --- | --- | --- | --- |
| Arabinogalactan | Gal tp | 0.257 | Galactose | Goellner et al. (59) |
|  | Gal 3p | 0.049 | Galactose |  |
|  | Gal 6p | 0.258 | Galactose |  |
|  | Gal 3,4p | 0.01 | Galactose |  |
|  | Gal 3,6p | 0.289 | Galactose |  |
|  | Ara tf | 0.045 | Arabinose |  |
|  | Ara tp | 0.048 | Arabinose |  |
|  | Ara 3f | 0.044 | Arabinose |  |
| Starch - Amylose (25%) | α-(1-4) | 0.25 | Glucose | Ai and Jane (66) |
| Starch - Amylopectin (75%) | α-(1-4) | 0.7125 | Glucose | Ai and Jane (66) |
| | α-(1-6) | 0.0375 | Glucose | |
| Inulin | $\beta$(2—1) | .862 | Fructose | Yao et al. (27) |
| | α-(1-2) | .138 | Glucose | |
| Cellobiose | $\beta$(1—4) | 1 | Glucose | French (68) |
| Maltose | α-(1-4) | 1 | Glucose | Nawaz et al. (67) |
| Fructans | Fru tf | 0.005 | Fructose | King et al. (69) |
| | Fru $\beta$(2—1)f | 0.045 | Fructose | |
| | Fru $\beta$(2—6)f | 0.34 | Fructose | |
| | Fru $\beta$(2—6,1)f | 0.22 | Fructose | |
|  | Glc tp | 0.05 | Glucose |  |
|  | Glc (1-6)p | 0.34 | Glucose |  |

The primary inputs for CheSL include the fractional composition of linkages in each carbohydrate and the fractional composition of carbohydrates in each medium. This information can be effectively represented by defining a linkage matrix (**L**) and media matrix (**M**), respectively.

The matrix **L**, representing the fractional composition of the linkage arrangements present across the carbohydrates contained in all media, can be given as follows:

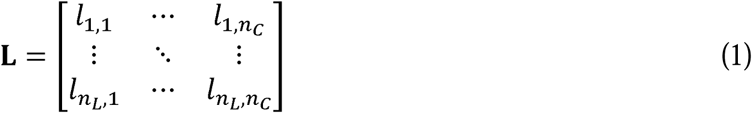

Where the rows and columns in **L** represent linkage arrangements and carbohydrates, respectively, *n_L_* denotes the total number of unique linkages associated with subunits contained Where the rows and columns in **L** represent linkage arrangements and carbohydrates, in all carbohydrates; the (*i,j*)*^th^* element of **L**, i.e., *l_i,j_*, denotes the proportion of linkage arrangement *i* within carbohydrate *j*.

The matrix **M** representing media-specific carbohydrate composition can be given as follows:

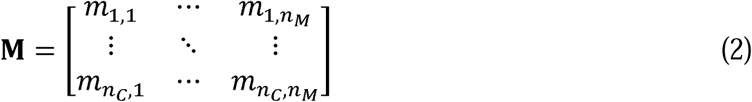

Where the rows and columns in **M** represent carbohydrates and media, respectively, *n_c_* and *n_M_* denote the total numbers of unique carbohydrates and media used in experiments, respectively; the (*j,k*)*^th^* element of **M**, i.e., *m_j,k_*, denotes the concentration of a specific carbohydrate *j* in a medium *k*.

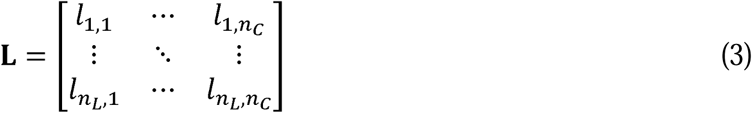

We can also define a (*n_L_* x *n_M_*) matrix **S** to represent the concentrations of linkage arrangements (or subunits) in media, which are readily obtainable by multiplying **L** and **M**, i.e.,

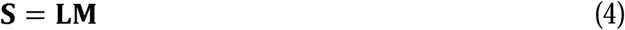

where the rows and columns in **S** represent linkage arrangements and media, respectively; the (*i,j*)*^th^* element of **S**, i.e., *s_i,k_*, denotes the *i^th^* linkage arrangement (or free subunit) concentration for the *k^th^* medium. The fractional abundance of the *i^th^* linkage arrangement (or free subunit) within the *k^th^* medium, denoted by *f_i,k_*, is simply calculated from *s_i,k_*(*i =* 1,2*, …, n_L_*) as follows:

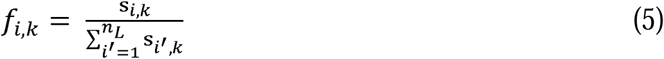

We can finally calculate CheSL Shannon diversity and CheSL Shannon evenness for the *k^th^*medium, denoted *CSI_k_* and *CSE_k_*, respectively, using the equations below:

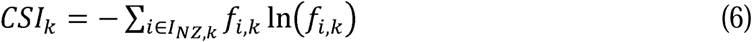

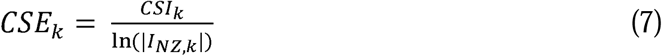

*I_NZ,k_* denotes the indices of non-zero elements of **s_k_**. For, CheSL Shannon evenness, we divide *CSI_k_* by the number of non-zero elements in **s_k_** (or the cardinality of *I_NZ,k_*, which represents the CheSL richness.

In the metric we developed, monomeric glucose was assigned a CheSL value of 0, because pathways to import and metabolize glucose are conserved across many species of bacteria (34–38) and as a result glucose metabolism was unlikely to provide a unique niche for microbes within most communities. Other unique monosaccharides (e.g., arabinose, N-acetylglucosamine, D-glucuronic acid) were assigned CheSL richness values of one, as pathways for their uptake and utilization were not expected to be as widely conserved as glucose and could represent unique niches. Other CheSL values for polysaccharides were primarily assigned based upon the number of unique linkage arrangements, as the capability to metabolize monosaccharides liberated from polysaccharides was expected to be part of the same unique niche. With the exception of soluble starch, carbohydrate composition and linkage data were estimated based upon previously published literature (Table 1).

Using the two hypothetical media exemplified in the bottom of Fig. 2, we illustrate how to calculate CheSL richness, CheSL Shannon diversity, and CheSL evenness metrics. Across these two media, there are three carbohydrates. Carbohydrate one has two linkage arrangements, of which 66% are arrangement one and 33% are arrangement two. Carbohydrate two was a polysaccharide, with three linkage arrangements (three, four, and five) in equal proportions. Carbohydrate three contains three linkage arrangements, 50% are linkage arrangement six, 33.33% are linkage arrangement seven, and 16.66% are linkage arrangement eight. The linkage matrix **L** is below:

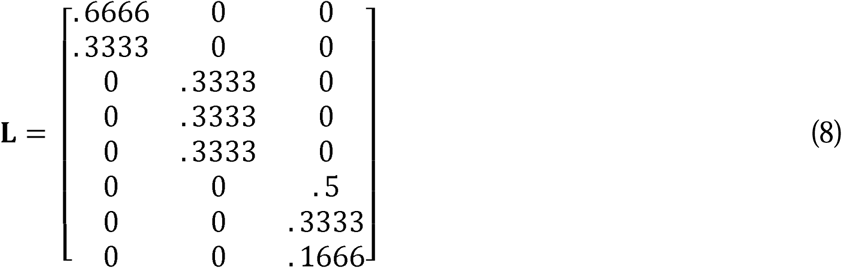

**Figure 2:**
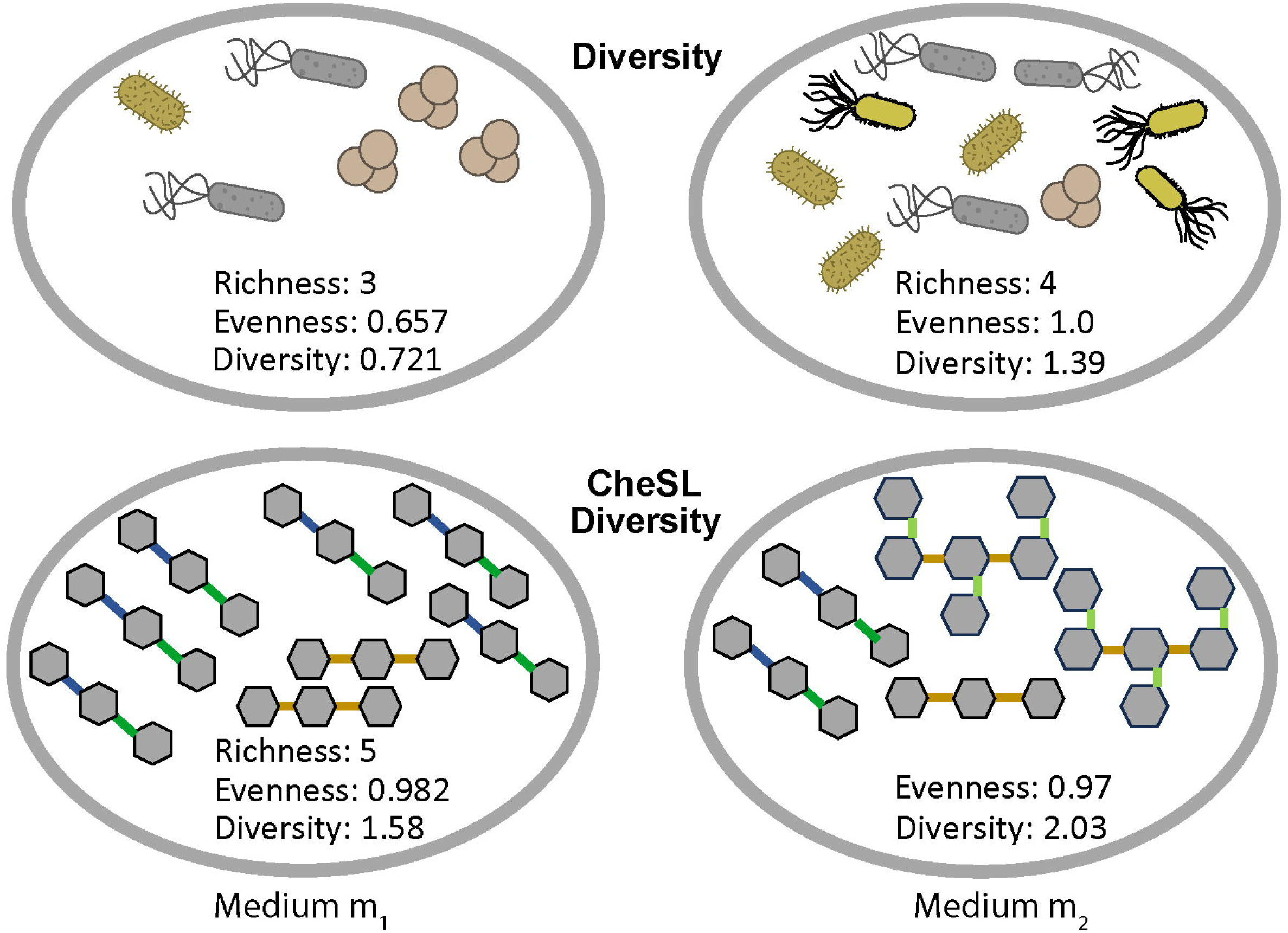
Graphical representation of Carbohydrate Chemical Subunits and Linkages (CheSL) metrics. Graphical representations of approaches to calculate microbial and CheSL richness, Shannon evenness and Shannon diversity. In these examples, different microbes are represented by different shapes and colors and different linkages are color-coded.

The first medium in the study, **m_1_**, contains 1 g/L carbohydrate one and 2 g/L carbohydrate 2. The second medium in the study, **m_2_**, contains 1 g/L carbohydrate one, 2 g/L carbohydrate two and 2 g/L carbohydrate three. The media matrix (**M**) is shown below:

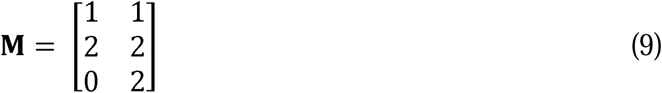

The concentration of linkages, **S**, can be calculated from L and M.

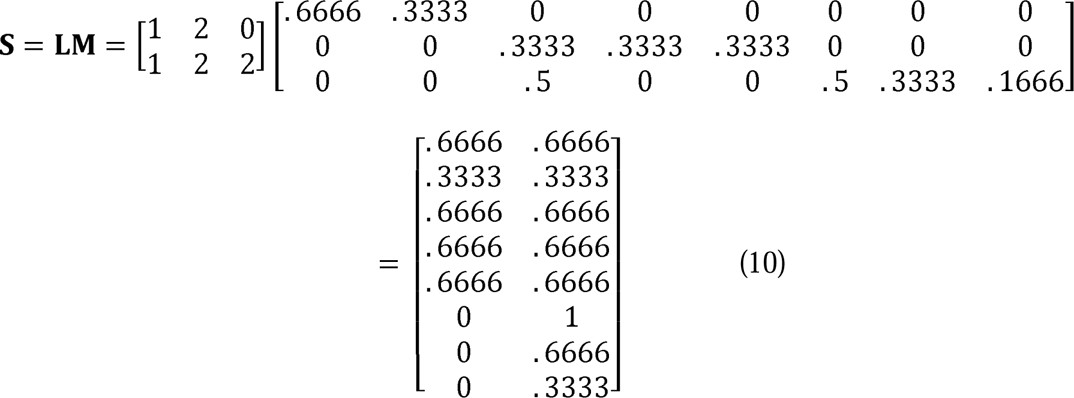

These calculations demonstrate that the medium, **m_1_** has 0.3333 g/L worth of carbohydrate associated with linkage two and 0.6666 g/L associated with linkage arrangements one, three, four, and five. **m_2_**has 0.666 g/L worth of carbohydrate associated with linkage arrangements one, three, four, five, and seven, 0.3333 g/L worth of carbohydrate associated with linkage arrangements two and eight, and 1 g/L worth of carbohydrate associated with linkage arrangement six.

From this, we calculated the fractional contributions of each linkage pool (f_j_ and f_k_). For f_j_, this was 0.222 for linkage arrangement one, three, four, and five, and 0.111 for linkage arrangement two. For f_k_, this was 0.1333 for linkage one, three, four, five, seven, 0.066 for linkage arrangements two and eight, and 0.2 for linkage arrangement six.

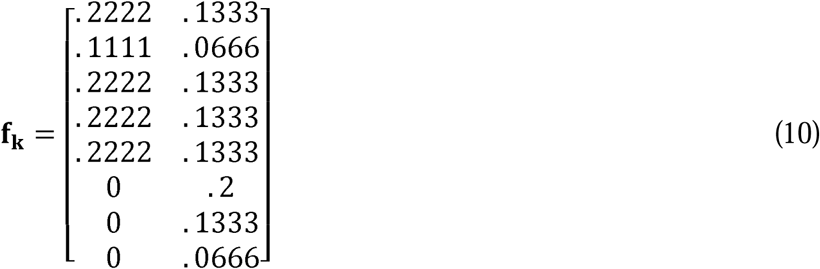

Using this fractional matrix and the formulas described above, we determined that for medium **m_1_,** the CheSL richness was 5, CheSL Shannon diversity was 1.58, and CheSL Shannon evenness was 0.982, whereas for medium **m_2_**, the CheSL richness was 8, CheSL Shannon diversity was 2.03, and CheSL Shannon evenness was 0.97.

### CheSL Shannon Diversity Connects Carbohydrate Complexity to Microbial Richness

We hypothesized that increasing the number of unique utilizable substrates would expand the fundamental niche space and facilitate increased microbial richness. To test this, microbial communities from the fecal samples of two healthy humans (Fecal Samples A and B (FSA, FSB)) were cultured in continuous-flow minibioreactor arrays (MBRAs) with media that differed in composition and abundance of carbohydrates. Previously, we demonstrated that community assembly in MBRAs was a result of environmental filtering that selected for core, cultivatable taxa as well as stochastic processes that contributed to variation between replicate communities (30). We expected that unique utilizable carbohydrate linkages would contribute to community assembly primarily through environmental filtering but that stochastic processes would also impact assembly.

Replicate cultures were inoculated into four media (NS, MM, 5MM, and IL) that varied in composition and abundance of carbohydrates but were composed of the same base of proteins, lipids, and trace minerals (**Table S1**). All media contained low levels of inulin and arabinogalactan. NS also contained low levels of mono (glucose) and disaccharides (cellobiose and maltose). MM contained low levels of mucosal monosaccharides and soluble starch. 5MM contained higher levels of mucosal monosaccharides and soluble starch compared to MM. IL was supplemented with ileostomy effluent, which likely contains a complex mixture of carbohydrates and other factors, requiring approximation of substrate concentrations from available literature (39, 40). Samples were collected from communities daily and changes in microbial community composition were determined through sequencing the V4 region of the 16S rRNA gene. Contamination limited collection of data from 5MM to FSB only.

Contrary to our original expectation, we found that microbiota richness (number of observed ASVs) was most strongly correlated with CheSL Shannon diversity rather than CheSL richness (Fig. 3). Specifically, we observed a Pearson correlation coefficient between ASV richness and CheSL richness of R=0.64 (p=0.0024, Fig. 3A), between ASV richness and CheSL Shannon evenness of 0.21 (p=0.38, Fig. 3B), and between ASV richness and CheSL Shannon diversity of R=0.74 (p=0.00019, Fig. 3C). Individual analysis of each fecal sample (Fig. S1A-F), indicated that the strong correlation was primarily a result of observations with FSB, as correlations for FSA were lower, which may have been a result of the relatively narrow range of CheSL Shannon diversity and CheSL Shannon evenness tested for FSA.

**Figure 3:**
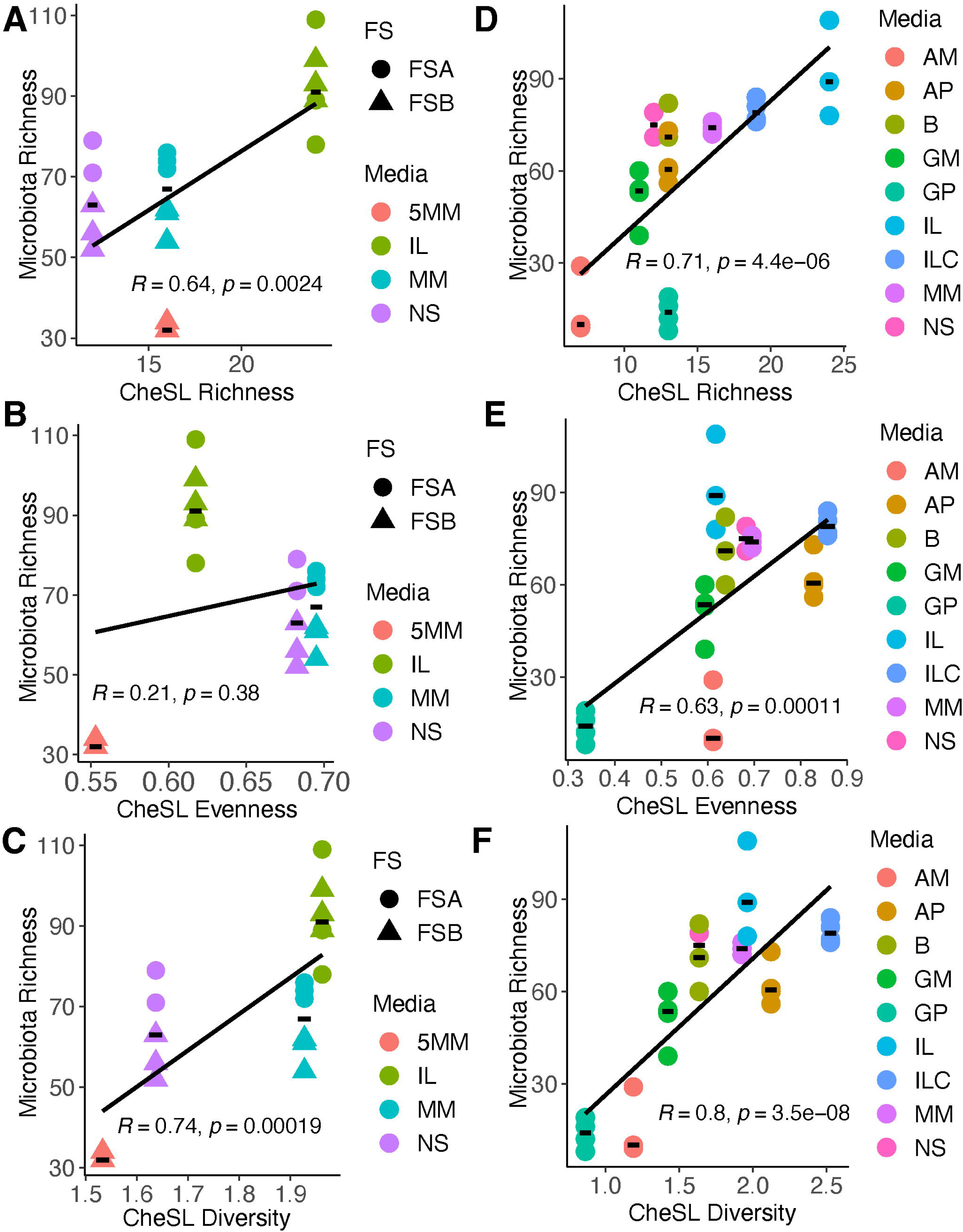
Effects of CheSL richness, CheSL Shannon evenness, and CheSL Shannon diversity on microbiota richness. Correlations of shared microbiota richness (observed ASVs shared across a time window as described in Methods) with (A) CheSL richness, (B) CheSL Shannon evenness and (C) CheSL Shannon diversity for FSA (circles) and FSB (diamonds) Correlations of shared microbiota richness with (D) CheSL richness, (E) CheSL Shannon evenness, and (F) CheSL Shannon diversity for FSA communities cultured in additional media along with those described in panels (A)-(C). In all plots, Pearson correlations coefficients and p-values are reported. Correlation analysis for FSA and FSB alone in the media shown in panels (A)-(C) and for FSA in the new media shown in panels (D)-(F) can be found in Supplementary Figure S1.

We evaluated the effects on microbial richness when FSA was cultured in additional media that varied across a broader range of CheSL Shannon Diversity and CheSL Evenness (Table 2). Unfortunately, FSB samples were no longer available for further testing. Medium B was similar to NS, but also had low levels of soluble starch. ILC included a mixture of carbohydrates inferred to be present in IL present at low concentrations (39, 40). The remaining four media had a single carbohydrate polymer or its monomers present at higher levels, but this carbohydrate varied in complexity. GP medium had higher concentrations of glucose polysaccharide (starch), GM medium had higher concentrations of glucose monosaccharide, AP medium had higher concentrations of arabinogalactan polysaccharide, and AM medium had high concentrations of the arabinogalactan monomers galactose and arabinose. We cultured replicate communities from FSA in these different media in continuous-flow MBRAs and analyzed changes in microbial community composition through sequencing of the V4 region of the 16S rRNA gene.

**Table 2:** Variations in Bioreactor Medium.

| **Carbohydrate (g/L)** | **B<sup>1</sup>** | **ILC<sup>1</sup>** | **GP<sup>1</sup>** | **GM<sup>1</sup>** | **AP<sup>1</sup>** | **AM<sup>1</sup>** |
| --- | --- | --- | --- | --- | --- | --- |
| Arabinogalactan | 0.1 | 0.05 | 0.1 | 0.1 | 2.0 | - |
| Inulin | 0.2 | 0.05 | 0.2 | 0.2 | 0.2 | 0.2 |
| Soluble starch | 0.2 | 0.05 | 2.0 | 0.2 | 0.2 | 0.2 |
| Cellobiose | 0.15 | 0.05 | 0.15 | 0.15 | 0.15 | 0.15 |
| Maltose | 0.15 | - | 0.15 | 0.15 | 0.15 | 0.15 |
| Glucose | 0.04 | - | 0.04 | 2.0 | 0.04 | 0.04 |
| Fructose | - | 0.05 | - | - | - | - |
| Galactose | - | 0.05 | - | - | - | 1.71 |
| Arabinose | - | - | - | - | - | 0.29 |
| N-acetylneuraminic acid | - | 0.05 | - | - | - | - |
| N-acetylglucosamine | - | 0.05 | - | - | - | - |
| L-fucose | - | 0.05 | - | - | - | - |
| D-glucuronic acid | - | 0.05 | - | - | - | - |
| Total carbohydrate (g/L) | 0.84 | 0.5 | 2.64 | 2.8 | 2.74 | 2.74 |

<sup>1</sup>Abbreviations for basal media variations: B: basal medium; ILC: ileostomy carbohydrates; GP: glucose polymer (soluble starch); GM: glucose monomer; AP: arabinogalactan polymer; AM: arabinogalactan monomers.

Adding this new data to our existing FSA data, we observed higher correlations between ASV richness and CheSL richness (Fig. 3D; R=0.71, p=4.4X10^−6^), CheSL Shannon evenness (Fig. 3E, R=0.63, p=0.00011), and CheSL Shannon diversity (Fig. 3F, R=0.8, p=3.5X10^−8^) across FSA samples. Analysis of FSA samples from this second experiment alone yielded higher correlations (Fig. S1G-S1I), more consistent with what was observed for FSB cultured across a range of media (Fig. S1A-S1C). As the two FSA experiments were performed on a cryopreserved fecal sample stored for years between experiments, the high correlation across the two experiments indicated the robustness of the observations.

We also evaluated two previously published data sets examining effects of carbohydrate diversity on fecal microbial community assembly (26, 27). For these data sets, carbohydrate linkage information was estimated from the available literature (Supplementary Table S2), and microbial richness data was obtained from the text and/or supplementary information. (26–28, 41–43). We observed similar correlations between microbiota richness and CheSL richness, CheSL Shannon evenness, and CheSL Shannon diversity (Fig. S2), across these studies, although substrate complexity and the needed approximations made for certain polysaccharides used (i.e., pectin), may have reduced statistical significance of data analyzed from Chung et al (26). Overall, these analyses demonstrated that CheSL Shannon diversity can be broadly applied, but also highlighted limitations for use with certain fecal communities and carbohydrate structures.

### CheSL Shannon diversity predicts peptide utilization

We used PICRUSt2 to infer genome content from 16S rRNA gene sequences and STAMP to identify differentially abundant functional pathways between communities cultured in different media. We focused on data from FSA cultured in the media described in Table 2 because we also had access to cell-free culture supernatants for further testing. One observation of interest, given our recent studies indicating Stickland fermentation contributes to persistence of *C. difficile* in fecal communities cultured in NS medium (44), was enrichment for taxa potentially encoding genes for L-arginine degradation (Stickland Fermentation; Fig. 4A) in microbial communities with higher shared richness (cultured in B, GM, AP, and ILC). We also observed predicted pathways for fermentation of pyruvate to butanoate (butyrate; Fig. 4B) were enriched in these same microbial communities. We hypothesized that differential abundance of Stickland fermentation pathways could indicate differences in peptide fermentation between communities. We found that peptide utilization varied between communities cultured in different types of media (Fig. 4C). When we performed a correlation between peptide utilization and CheSL Shannon diversity, we observed a strong correlation (R = −0.77, P = 9.5X10^−6^, Fig. 4D), similar to that observed between peptide utilization and microbiota richness alone (Fig. S3A) and higher than that observed for CheSL Shannon evenness (Fig. S3B). From this data, we infer that for communities cultured in the presence of more complex mixtures of carbohydrates, peptide utilization may serve as an additional substrate niche dependent on community composition; alternatively, these communities may have higher needs for peptide utilization to facilitate carbohydrate degradation capabilities.

**Figure 4.**
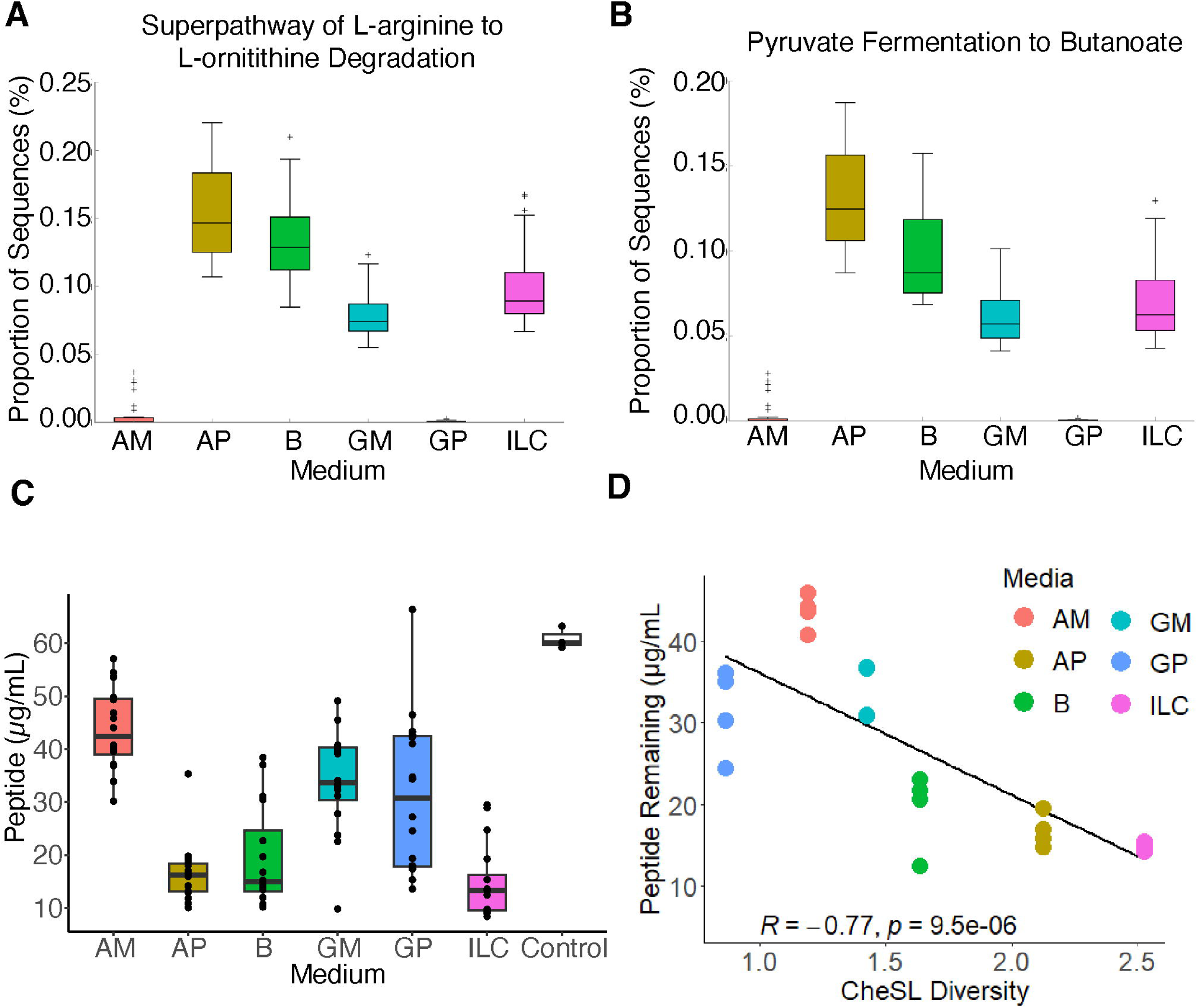
Differentially abundant functional pathways suggest divergent metabolism and production of beneficial compounds. (A-B) STAMP generated plots of pathways differentially abundant in FSA communities cultured in media with high CheSL Shannon diversity complexity and low CheSL Shannon diversity. (C) Total peptide concentrations measured in steady state culture of each media type exhibiting differences in peptide utilization by communities compared to fresh media control (D) Correlation between CheSL Shannon diversity and the peptide concentration in spent medium. Pearson correlation coefficient and p-value are reported. Additional correlations can be found in Supplementary Figure S3.

### *Bacteroides* and *Lachnospiraceae* are central members in conserved, rich communities

As microbial interactions can have significant impacts on microbial community assembly, stability and function (25), we investigated how variations in carbohydrate composition affected microbial interactions. We used LIMITS, a computational algorithm that infers microbial interactions by fitting a discrete Lotka-Volterra model against the temporal profiles of ASV relative abundance data (45, 46), to analyze our sequence data. The LIMITS equation was derived by implicitly assuming that total community biomass does not significantly vary in time. Thus, performance is improved when LIMITS is conducted with data collected from communities with minimal variations of the total biomass, such as those found when culturing microbial communities under continuous-flow conditions approaching steady state. However, LIMITS ignores temporal variations of interspecies interactions by assuming interaction coefficients to be constant in time. While relevant in certain cases, this assumption is invalid in general (47, 48). To address this limitation, we employed a moving window approach by: (1) dividing the data into multiple subsets, each representing a shorter time period; (2) using LIMITS to identify (constant) microbial interactions within these shorter periods; and (3) iteratively inferring the network by shifting the time window to subsequent timepoints. By analyzing the goodness of fit of these interaction networks (mean cross-correlation between inferred networks and observed networks) across time windows, we observed that the mean cross-correlation across media types increased after the early phase of *in vitro* culture (Fig. S4).

We initially compared ensemble networks at the ASV level across replicate communities cultured in the same media. However, very few interactions were conserved, which is consistent with lower levels of ASV conservation observed between replicate communities. We posited that neutral processes acting during initial community assembly likely contributed to variations in ASV composition but that environmental filtering due to carbohydrate composition could select for ASVs with similar functions. We assessed conserved interactions at the genus level, as conserved functions have been previously observed to occur at the genus level in human fecal communities (49), and identified interactions conserved between replicates within a medium.

Our analysis of the ensemble models primarily focused on FSA samples cultured in B, AP, AM, GP, GM and ILC (Fig. 5), because the frequency of sample collection in this study facilitated higher quality ensemble models. (Data from FSA and FSB in NS, MM, and IL can be found in Fig. S5). We saw a distinct trend in the number of nodes, representing interacting taxonomic groups, and the shared richness of the cultures. (Fig. 5A). These networks were not driven solely by correlations between genera, but rather population dynamics fit to generalized Lotka Volterra models by LIMITS (45), as was demonstrated by different network features between media (e.g., AP, ILC, and B). However, it is important to note that not all taxonomic groups were represented in the networks, as not all groups had conserved (or any) interactions detected in some conditions.

**Figure 5:**
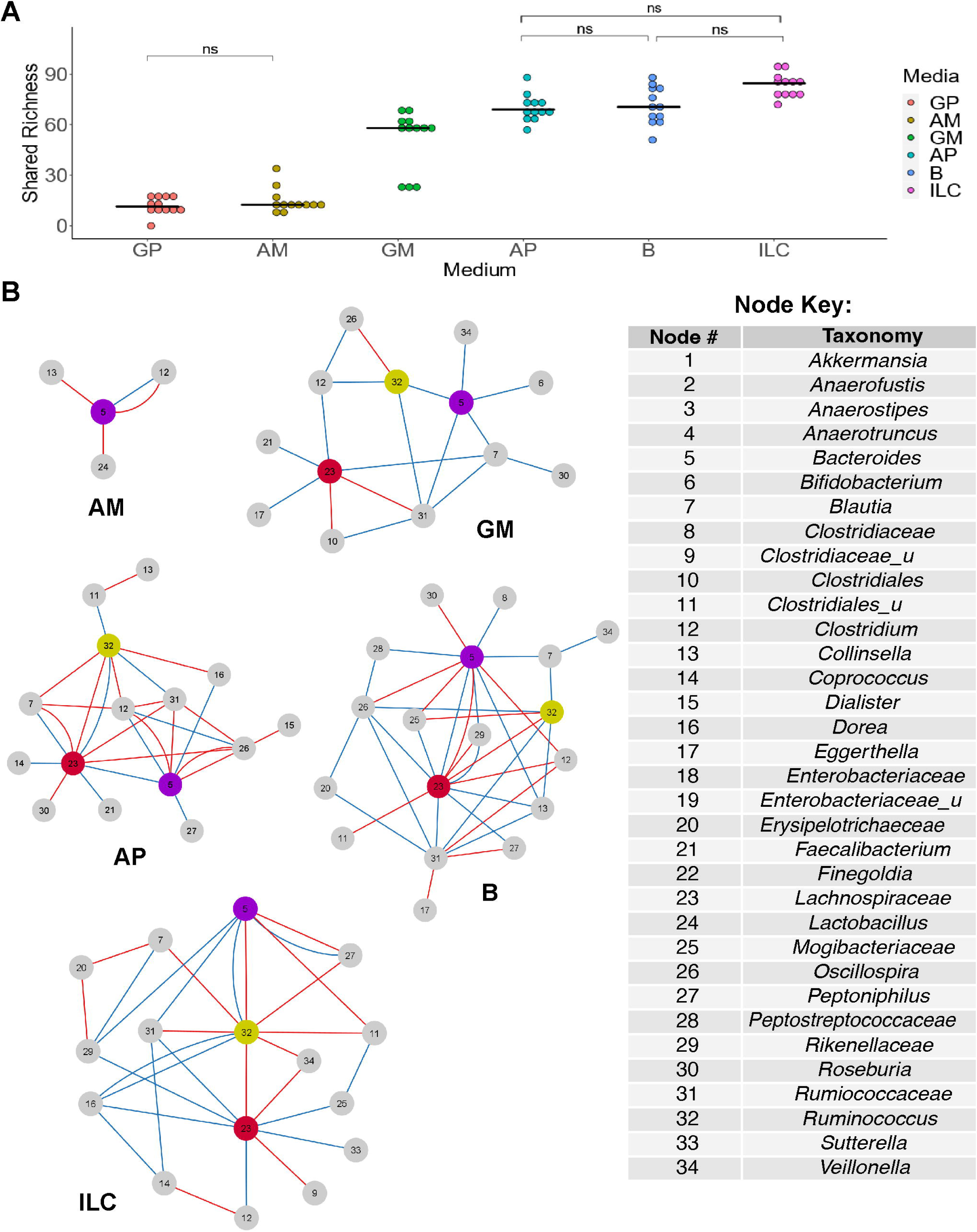
Ensemble Networks of Conserved Interactions demonstrate conserved structure of communities cultured in media that support high shared microbiota richness and within-group similarity. (A) Shared microbiota abundance measured across three late culture windows for each FSA replicate culture (n=4), cultured in the media indicated. Unless otherwise noted (ns, for non-significant) comparisons between medium were highly significant with an adjusted p-value < 5×10^−5^ from Tukey’s HSD (B) Conserved ensemble networks were built for each media from the networks inferred from the window with the highest quality of fit for each replicate cultured in AM, GM, AP, B and ILC as marked below each network. A key describing taxa for each numbered node is provided. Positive interactions are indicated by red lines and negative interactions are indicated by blue lines. An ensemble network for GP could not be determined due to lack of conserved interactions.

We found three consistently central taxonomic groups, *Bacteroides*, *Lachnospiraceae*, and *Ruminococcaceae* of unclassified genera (Fig. 5B). Particularly *Lachnospiraceae* and *Ruminococcacecae* of unclassified genera exhibit conserved centrality in media with high CheSL Shannon Diversity which produce complex ensemble networks. While the loss of *Lachnospiraceae* spp. interactions in low CheSL Shannon Diversity networks could possibly be explained by differences in abundance between media types (e.g., the negative log_2_ fold change in GP compared to B), differences in abundance of *Ruminococcaeceae* spp. could not explain the absence of interactions between communities cultured in different media, as these species were not differentially enriched (Fig. 6). We also found greater connectivity in conserved networks among media types which supported the highest microbiota richness. To support future uses of ILC medium, we found communities cultured in this medium exhibited more conserved, positive interactions between the three central taxonomic groups mentioned earlier (Fig. 5F). Overall, we observed more conserved relationships across replicate cultures with higher CheSL Shannon diversity, suggesting that this metric could be used to not only increase microbiota richness, but also the reproducibility of microbial interactions in cultures derived from complex sources.

**Figure 6:**
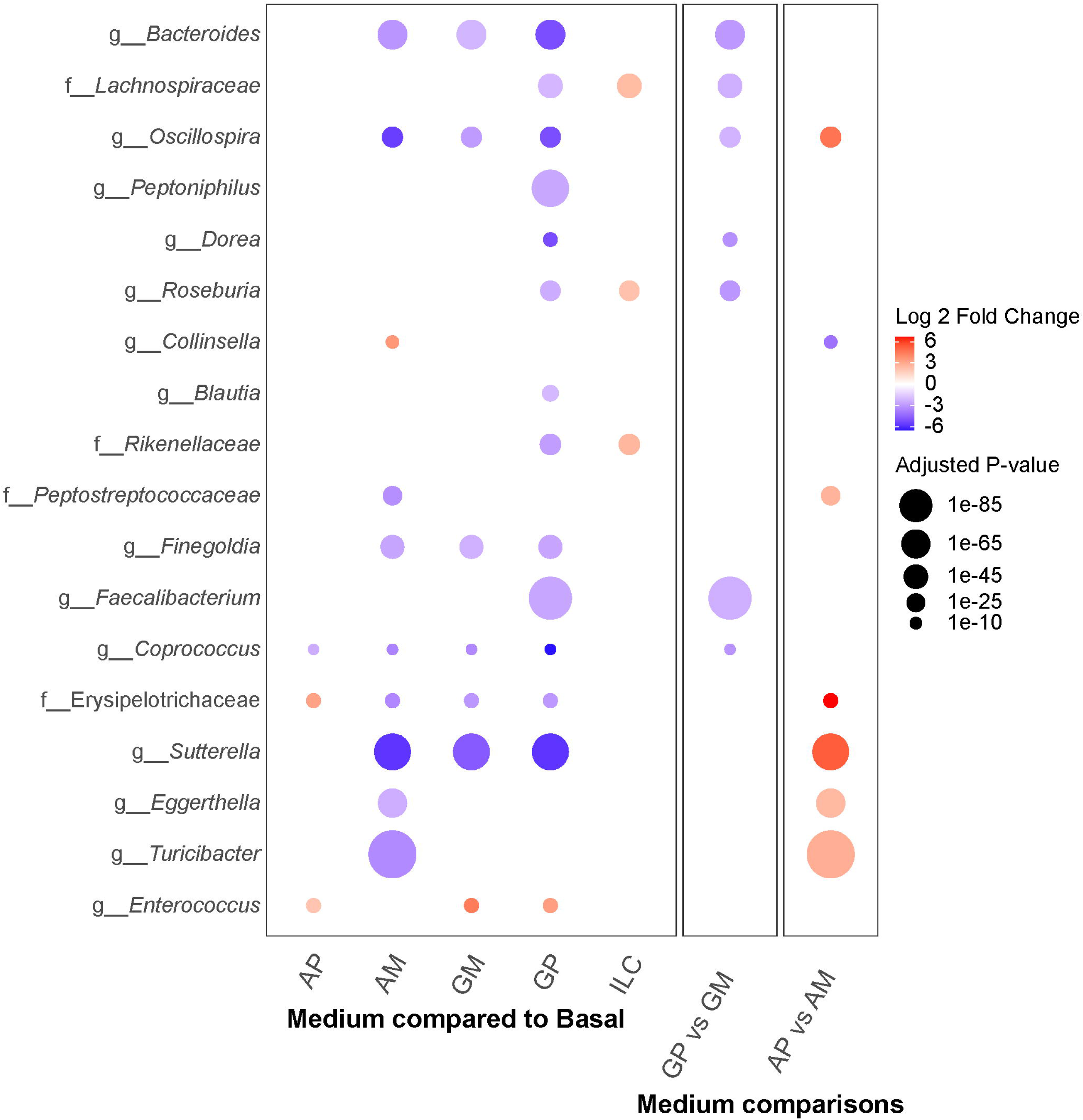
Differential Abundance of Taxa Between FSA Communities Cultured in Different Media. ASVs were binned into shared taxa at the genus level (or lowest level of taxonomic resolution available for taxa whose classification could not be determined at the genus level). Differential abundance of taxa between communities cultured in different media types was determined using DESeq2. Color indicates the log_2_-fold change in abundance between cultures and the size of the points indicate the statistical significance of the abundance difference. Unless otherwise indicated, media were compared to medium B as baseline. Comparisons were also made between communities cultured in overrepresented polysaccharide media (GP and AP) and their respective overrepresented monosaccharide media (GM and AM).

For media with low conservation of interactions between replicate networks (i.e., GP and AP), we gained some insights into the effects of media on community composition through differential enrichment analysis (Fig. 6). We observed that most taxa decreased in abundance in GP compared to B (Fig 6), except for *Clostridium* spp. and *Bifidobacterium* spp (Supplementary Table 3). For AM, we saw a similar selection against many genera, but enrichment of *Collinsella* spp. (Fig 6). Based upon the ensemble network, this enrichment may be driven by *Bacteroides* spp. (Fig. 5B, 6). Alternatively, enrichment may be due to the comparatively low ratio of monosaccharides to polysaccharides, as was previously found in human fecal samples (50).

## DISCUSSION

In this work, we developed novel metrics to describe carbohydrate complexity based on Chemical Substrate and Linkage (CheSL) richness, CheSL Shannon diversity, and CheSL Shannon evenness and demonstrated that mixtures of carbohydrates with higher levels of CheSL Shannon diversity supported growth of human fecal microbial communities with higher richness and conserved microbial interactions. This was contrary to our initial hypothesis that CheSL richness would be the primary driver of microbial richness, and points to a role for CheSL Shannon evenness (which also contributes to CheSL Shannon diversity) in microbial community assembly. We hypothesize that CheSL Shannon evenness is most likely to contribute to increased microbiota richness when carbohydrate mixtures are highly uneven. Specifically, when species who occupy different carbohydrate niches compete for shared resources, generation of higher levels of biomass by the species who occupy the carbohydrate niche with higher concentrations may deplete other limiting nutrients that will lead to the extinction of other species unable to compete for this limiting nutrient. Thus, these observations do not contradict, but rather build upon nutrient-niche theory (23, 24) and integrate the concepts of competitive fitness in the context of community stability and carbohydrate diversity.

This work builds on the body of emerging literature linking carbohydrate complexity to microbial community composition and function. Similar to what was observed by Chung et al. and Yao et al., media with higher levels of CheSL Shannon diversity support higher levels of microbiota richness in communities cultured from fecal samples across the samples tested (26, 27). As Loss et al used a defined community of 10 microbes selected to provide representation of microbiota diversity, they did not see effects of carbohydrate complexity on richness (29). However, reanalysis of their data demonstrated correlations between microbiota diversity and carbohydrate complexity (Fig. S6). We also observe correlation between CheSL Shannon Diversity and microbial diversity (Fig. S6), but the correlations observed with microbiota richness were of primary interest for this study. Of note, the CheSL Shannon Evenness metric that we describe is similar to the carbohydrate complexity measure determined by Loss (29). Our work expands on their studies through development of CheSL Shannon diversity, which for the data in this study, exhibits stronger correlation with microbiota richness than CheSL Shannon evenness.

Other studies have found more complex relationships between carbohydrate complexity and microbiota richness that are likely due to differences in ability of microbial communities to cleave linkages identified (28, 51, 52). Degradation of complex polysaccharides is dependent upon the presence of specific carbohydrate active enzymes (CAZymes) within microbial community members (53). In the absence of enzymes able to break unique linkages within complex polysaccharides, monosaccharides remain inaccessible and cannot provide unique niches. The presence of specific CAZymes which allow degradation of linkages are the gateway to energy for the microbes, and in turn, linkages serve as a representation of the latent energy granting niche space. Thus, it was not entirely surprising when we observed that when more carbohydrate mixtures and microbial communities were analyzed, correlations between CheSL Shannon diversity and microbiota richness were lower (Fig. S2). However, we suggest that predictive accuracy of CheSL can be improved with prior knowledge of the CAZyme diversity within the complex community being targeted. Inherent assumptions used in this study were that the complex fecal microbiota would be capable of utilizing the selected carbohydrates as they are abundant in diet or are host derived. One should be careful in use of CheSL for predictions of in cases of low initial functional diversity or addition of uncommon substrates to communities which likely lack the enzymatic potential for degradation (e.g., cellulose to human fecal sample derived community). Further, the CheSL metrics should be applied across media which are controlled with respect to other substrates. Finally, it is also important to note that CheSL measures are estimates based upon published literature and will improve as additional carbohydrate composition and linkage information is published

From our observations, among samples cultured in media with the greatest CheSL Shannon diversity (B, AP, and ILC), there is higher similarity between microbial interaction network structure and shared microbiota richness. This may suggest the diversity of nutrient niches inferred by the abundance and evenness of linkages may control the reproducibility of community assembly from complex microbial samples in continuous flow bioreactors, thereby impacting the shared microbiota richness of the samples. Loss and colleagues similarly observed that higher carbohydrate complexity led to higher reproducibility in community assembly (29). From our findings, the initial assembly of a community around *Bacteroides*, *Lachnospriaceae,* and *Ruminococcaceae* resulted in a network of interactions supporting reproducible cultures with greater microbiota diversity under these culture conditions.

We also found enrichment and microbial associations previously observed in *in vivo* systems exposed to similar treatments– another indicator of our *in vitro* system’s capability to support representative communities. Particularly the relationship between *Bacteroides* and *Lachnospiraeceae* were supported through studies demonstrating cross feeding between *Bacteroides thetaiotaomicron* and *Anaerostipes caccae* (54)*. Clostridium sensu stricto* and *Bifidobacterium* has previously been observed in humans consuming diets high in starch aligning with our GP medium - which over-represents soluble starch for which these cultures continued to support of these taxa despite significant reduction in diversity (31, 55, 56).

Communities formed in conditions with high CheSL Shannon Diversity also exhibited enrichment of genes for potential Stickland fermentation of amino acids. Consistent with these observations, we observed higher levels of peptide depletion in these media. Future studies will be needed to determine whether utilization of peptides in media with high CheSL Shannon Diversity is conserved across other microbial communities and media types. Typically, proteolytic fermentation is associated with negative health outcomes, with a tradeoff between saccharolytic and proteolytic microbes (57). In our studies, peptide consumption occurred in the presence of dietary fibers and in communities enriched with genes for butyrate production, suggesting that potential beneficial functions of the microbiota were also maintained.

Like any ecological set of interactions, interactions between GI microbiota are directly determined by the biotic and abiotic properties of the environment to which the community is exposed. Using approaches that adhere to limitations in data analysis and modeling as described in this work, we gained additional resolution into the relationships between fecal bacteria and the potential interaction landscapes which can be realized from the availability of specific substrates. From progression in process-based approaches to interaction prediction, we further informed and connected population dynamics with metabolic potential and ecological theories of nutrient niche partitioning (23, 24). Using CheSL Shannon diversity as a guide when designing *in vitro* culture media has the potential to improve reproducibility and increase the similarity of *in vitro* communities to those found in the fecal samples without introducing unknown dietary components that would be found in media supplements, such as ileostomy effluents or foods that have undergone simulated digestion. Our initial test at an improved carbohydrate medium supports this potential, with ILC granting a similar network structure to IL and the greatest shared richness of media which had a known substrate concentration (IL being of complex origin). By improving the reproducibility of community assembly in continuous flow conditions, we can better detect differences between samples and reactors driven by other interventions. Additionally, the experimental conditions we used, the improved quality of fit realized using a continuous flow bioreactor, strict filtering, and division of our time series into windows which represent transitory states in the community, all together may aid in inferring interactions for future mechanistic investigation of microbial metabolic interactions.

## MATERIALS AND METHODS

### Samples used in this study

Fecal samples were previously collected from two adult volunteers who were self-described as healthy and had not consumed antibiotics within the previous two months (30). Samples were homogenized under anaerobic conditions, aliquoted into anaerobic tubes and stored at −80°C until cultivation. Ileostomy effluents were collected from participants who had ostomies placed due to previous illness or injury as previously described (58). Samples were stored at −20°C until use. Informed consent was obtained prior to sample collection according to a protocol approved by the Institutional Review Board of Michigan State University (10-736SM).

### Media used in this study

Microbes from fecal samples were cultured in bioreactor basal medium that varied in carbohydrate composition. 1 L of basal medium contained 1 g tryptone, 2 g proteose peptone #3, 2 g yeast extract, 0.4 g sodium chloride, 0.5 g bovine bile, 0.01 g magnesium sulfate, 0.01 g calcium chloride, and 2 mL Tween 80, which was sterilized by autoclaving. 2 g sodium bicarbonate, 0.2 g inulin from chicory, 0.1 g arabinogalactan from larch wood, 0.2 g soluble starch, 0.04 g potassium phosphate dibasic, 0.04 g potassium phosphate monobasic, and 0.001 g vitamin K3 were filter sterilized and added to each liter of basal medium after autoclaving. Different variations of carbohydrates added to basal medium were used to test the impact of carbohydrate composition and concentration on microbial communities (Table S1). NS (No starch) lacked soluble starch and contained 0.15 g/L D-cellobiose, 0.15 g/L maltose, and 0.04 g/L D-glucose in addition to basal medium. MM (mucosal monosaccharides) contained an additional 0.2 g/L of soluble starch, 0.05 g/L N-acetylglucosamine, 0.05 g/L N-acetylneuraminic acid, 0.05 g/L D-glucuronic acid, and 0.05 g/L L-fucose. 5MM (5X higher mucosal monosaccharides) contained an additional 1.8 g/L soluble starch (2g/L final concentration), 0.2 g/L N-acetylglucosamine, 0.2 g/L N-acetylneuraminic acid, 0.2 g/L D-glucuronic acid, and 0.2 g/L L-fucose (0.25 g/L final concentration each). GP (glucose polysaccharide) contained an additional 1.8 g/L starch (2 g/L final concentration). GM (glucose monomer) contained an additional 1.96 g/L D-glucose (2 g/L final concentration). AP (arabinogalactan polymer) contained an additional 1.9 g/L of arabinogalactan (2 g/L final concentration). AM (arabinogalactan monomers) lacked arabinogalactan and instead contained the two primary monosaccharides present in arabinogalactan (59, 60)- 1.79 g/L galactose and 0.21g/L arabinose. IL (ileostomy effluent) contained an additional 0.2 g/L soluble starch, 0.05 g/L N-acetylglucosamine, 0.05 g/L N-acetylneuraminic acid, 0.05 g/L D-glucuronic acid, 0.05 g/L L-fucose, and 100 mL of ileostomy effluent. Ileostomy effluent was thawed, pooled from multiple donors, and filtered through a coffee filter to remove particulates prior to sterilization along with base medium by autoclaving. Carbohydrate diversity of ileostomy effluent was estimated from references (39, 40). Finally, ILC (ileostomy carbohydrates) reduced the concentrations of arabinogalactan, inulin, and soluble starch to 0.05 g/L for each of these carbohydrates and also contained 0.05 g/L cellobiose, 0.05g/L fructose, 0.05g/L galactose, 0.05 g/L N-acetylneuraminic acid, 0.05 g/L N-acetylglucosamine, 0.05 g/L L-fucose, and 0.05 g/L D-glucuronic acid.

### Culturing human GI microbes in continuous flow bioreactors

We used MBRAs (30), continuous-flow culture vessels that simulate environmental parameters of the lumen of the distal human colon and maintain communities in steady-state based on the carrying capacity of the reactors, to culture communities of human GI microbes from fecal samples. MBRAs are parallelized, therefore, we could culture replicate communities of human GI microbes in medium that varied by carbohydrate composition and concentration and determine how these changes impacted microbial interactions. Prior to inoculation into MBRAs, fecal samples were thawed and homogenized in anaerobic phosphate buffered saline (PBS) as a 20% w/v slurry as previously described (30). In the first experiment, Fecal Sample A (FSA) was cultured in NS (n=2 replicates), MM (n=3 replicates), or IL (n=3 replicates) and Fecal sample B (FSB) was cultured in NS (n=3 replicates), MM (n=3 replicates), 5MM (n=3 replicates), or IL (n=3 replicates). Media contamination prevented cultivation of FSA in 5MM, whereas technical difficulties led to loss of third replicate of FSA in NS. Both FSA and FSB formerly referred to as Fecal Sample B and Fecal Sample C respectively, were previously characterized when cultured in BRM2, a medium highly similar to NS, with an 8 hour retention time (30). For these experiments, 16 hours after inoculation, continuous flow was initiated at 0.94 mL/hr (16 hr retention time for 15 ml minibioreactor). One mL samples were collected from each culture daily for 10 days as previously described (30).

In the second experiment, FSA was selected for studies because of fecal sample availability. FSA was cultured in B, GP, GM, AP, AM, and ILC media, with n=4 replicates/medium. As before, continuous flow was initiated 16 hr after inoculation at 0.94 mL/hr. 1 mL samples were collected every 12 hours for 8 days. The increased frequency of sampling and number of replicates was to improve our ability to subsequently infer microbial interactions from data. In both experiments, MBRAs were maintained in an anaerobic chamber (Coy Laboratories) heated to 37°C with a gas atmosphere of 90% N_2_, 5% CO_2_, and 5% H_2_.

### 16S rRNA gene amplicon sequencing

The V4 region of the 16S rRNA gene was amplified directly from mechanically disrupted cells with broad range 16S rRNA primers 515F/805R as previously described (30). Purified amplicons were pooled in equimolar concentration and sequenced using Illumina MiSeq v2 according to Manufacturer’s protocol. Sequence data were processed using mothur V1.35.0. Sequences were aligned using the Silva r132 16S rRNA gene sequence database (61). Any chimeric sequences were removed following identification by UCHIME, and any contaminating sequences identified as chloroplasts, mitochondria, Archaea or Eukarya are removed as well (62). Sequences were denoised using mothur’s pre.cluster function, giving ASVs, and taxonomy was assigned using the Silva r138 16S rRNA gene sequence database. Downstream analysis of the taxonomic and count data was handled in R, using the phyloseq package (63, 64); code will be published on Github upon acceptance. An ASV table is provided in Supplementary Table S3, which is posted at https://github.com/AuchtungLab/CheSL.

### Data pre-processing for computational network inference

Since Lotka-Volterra equations model niche-based effects determining population dynamics, multiple filtering steps were required to mitigate spurious interactions. During the early phases of culture, two factors could influence community stability: the loss of ASV presence as non-viable cells were removed by dilution and partnership switching as communities adapted to the culture conditions. To improve LIMITS inferred model quality of fit, we tested whether a sliding window approach, sampling time regions across the time series, could improve consistency of interactions over time. We tested windows of varying sizes from 6 to 15 time points, across the entire time series. To inhibit performance loss from transient ASVs, we filtered ASVs by minimum presence across all time points within a window, requiring presence of a single read from an ASV across all time points. This pre-processing avoided issues associated with taxa falling and rising above the threshold of detection, which would influence the model similarly to migration and extinction events, and disturb inferences of a fixed ecosystem’s interactions (65). From this filtering we obtained what we call a “shared microbiota richness” which represents the count of ASVs found consistently across time, excluding ASVs that may be alternating above and below threshold of detection.

### Inference of interspecies interactions using LIMITS

LIMITS analysis was performed using the seqtime package in R by Faust et al. (46). Prior to running LIMITS, we preprocessed the data as described above. From there, we applied the LIMITS algorithm using a sliding window approach as described above. Inferred interactions were then used to simulate time series for the given window of time and the resulting simulated data were compared to the experimental data. Mean cross correlations were used to quantify the quality of fit of the simulated data to the experimental data. Specifically, mean cross correlations were calculated by determining the correlation between each taxon’s simulated and experimental data over the time series; the mean correlation of all taxa was then calculated; this method of quality of fit analysis was also implemented through the seqtime package (46). This sliding window method granted greater quality of fit than applying LIMITS to the entire experiment’s time series. In addition to improved quality of fit, a sliding window approach allowed us to select the time windows which most likely represent interactions driven by deterministic effects, rather than including data from the first day(s) of cultivation which likely included influences of founder effects, partner selection, and adjustment to the bioreactor system.

From each replicate and window, a network was inferred using LIMITS. To identify conserved interactions, we calculated consensus across networks from replicates of the same fecal sample and media culture conditions. using the same fecal sample inoculum. After calculating average interaction values and filtering by a consensus threshold of 2, meaning at least 2 networks identified the interaction with the same directionality, we obtained ensemble networks for each media type.

### Characterization of chemical subunit and linkage diversity

As described in the results, we developed CheSL Richness, CheSL Shannon diversity and CheSL Shannon evenness measures to calculate the complexity of carbohydrate mixtures based upon distinct linkages within di- and polysaccharides or the presence of unique monosaccharides. For example, the polysaccharide arabinogalactan is primarily composed of arabinose (80%) and galactose (15%) monomers (59) that are linked in eight unique types linkages within the polysaccharide (Table 1), with the relative abundances of these linkages are also known. We additionally informed the ratios of our linkages for amylopectin from literature, which indicated predominantly α-1–4 linkages, with 5% of linkages being α-1–6 linkages (66). For disaccharides, which have a single linkage type, there is no effective change in diversity when applying CheSL if it is the sole carbohydrate, but it is important to decompose into linkage types in cases where there is overlap. For example, we find that maltose, with α-1–4 linkages, contributes to the same pool as amylopectin (67). While cellobiose and inulin, have their own unique linkage pools - β(1—4) glucose for cellobiose and β(2—1) fructose and α-(l—2) glucose for inulin (27, 68). We estimated fructans linkage diversity from a study containing glycosidic linkage proportions from wheat kernels, yielding six distinct linkage types (69). For estimations of linkage diversity for CheSL calculations to apply to Chung et al.’s data, we referenced literature for pectin, beta-glucan, and glucomannan respectively (41–43). Composition and linkage data not described here can be found in Table 1 and in Supplementary Tables S1 and S2. Fig. 2 provides a representation of how CheSL diversity can be calculated across a medium containing multiple carbohydrates.

### Statistical analysis – comparison of means and correlations

To make comparisons for media shared richness, we used the R package MASS’s implementation of a negative binomial GLM, glm.nb, as the richness did not fit a normal distribution determined by the Shapiro-Wilk test, and the data was over-dispersed for a Poisson distribution (70). Pairwise comparisons were made with the R package emmeans and Tukey’s HSD was used to adjust for family-wise error rate (71). For correlations, the R Package ggpubr was used alongside tidyverse and ggplot2 to visualize and calculate the Pearson correlation between diversity metrics (72–74).

### Inferred metabolic functional potential with PICRUSt

After characterization of conserved interactions and visualization using Cytoscape, we inferred Enzyme Commission Pathway abundance for our cultures from fully sequenced isolates with similar 16S rRNA gene sequences using PICRUSt2 (75). The potential significance of Pfam pathway abundance was assessed using STAMP (76).

### Measurement of amylose/amylopectin ratio

To measure the ratio of amylose/amylopectin for this study we use a NEOGEN Megazyme amylose/amylopectin Assay kit, product code K-AMYL. The assay was performed according to the manufacturer’s instructions. This kit was used to determine the proportions of amylose and amylopectin comprising the soluble starch used in various media formulation. This ratio was then used to calculate CheSL diversity measures.

### Differential analysis of microbial abundances

We performed differential abundance analysis with DESeq2 (77) between communities cultured in basal bioreactor medium (B) and media with differing carbohydrate composition. The DESeq2 model we used to explain composition as a function of media type, with time as a covariate. Additionally, we performed DESeq2 analysis between communities cultured in media composed of the polymer form (starch (GP), arabinogalactan (AP)) and respective monomer forms (glucose (GM), galactose and arabinose (AM)), as they represent potential differences between monosaccharide-based niches and linkage-based niches at the genus level. Changes in abundance that were identified by DESeq2 as significant and had an absolute fold-change in abundance greater than two were plotted.

## Acknowledgements

The authors thank Dr. Robert A. Britton (Baylor College of Medicine, Houston, Texas USA) for access to sequence abundance data collected in his laboratory by JMA and shown in Fig. 3A-3C as well as access to the FSA fecal sample used in follow up experiments.

H.C.M and J.M.A. were responsible for conceptualization and design of the studies. Data collection was performed by H.C.M. and J.M.A. Analysis was performed by H.C.M. with guidance provided by H.-S.S. and J.M.A. J.M.A. acquired funding for the studies. The initial draft of the manuscript was written by H.C.M. and J.M.A. and all authors revised and edited the final manuscript. Bioinformatics was done using the Holland Computing Center, supported by the UNL Office of Research and Economic Development and the Nebraska Research Initiative.

## Data Availability

16S rRNA gene data is available in the NCBI Sequence Read Archive under BioProject ID PRJNA1146550.

## Ethics Approval

Protocols for collection and use of fecal samples were reviewed and approved by Institutional Review Boards at Michigan State University (10-736SM) and University of Nebraska-Lincoln (protocol number 18585).

## SUPPLEMENTAL MATERIAL

**Supplementary File 1: Figures S1 to S6**

**Supplementary Figure S1. Effects of CheSL richness, CheSL Shannon evenness, and CheSL Shannon diversity on microbiota richness within fecal sample communities and between experiments.** Correlations between shared ASV richness and (A,D,G) CheSL Richness, (B,E,H) CheSL Shannon evenness, and (C,F,I) CheSL Shannon diversity are shown FSB (A-C) and FSA (D-F) cultured in NS, IL, MM and 5MM (A-C only), and for FSA cultured in B, AM, AP, GM, GP, and ILC (G-I). Pearson correlation coefficients and p-values are reported.

**SupplementaryFigure S2. Effects of CheSL richness, CheSL Shannon evenness, and CheSL Shannon diversity on microbiota richness from previously collected data.** (A-C) Correlations between shared OTU Richness and (A) CheSL richness, (B) CheSL Shannon evenness, and (C) CheSL Shannon diversity from data reported by Yao and colleagues (27). (D-F) Correlations between shared OTU richness on days 15, 18, and 20 for Donor 1 (calculated from data published as Supplementary Table S1) and (D) CheSL richness, (E) CheSL Shannon evenness, and (F) CheSL Shannon diversity from data reported by Chung and colleagues (26). Supplementary Table S2 provide details for CheSL calculations. Pearson correlation coefficients and p-values are reported.

**Supplementary Figure S3. Reduction in peptide levels strongly correlates with microbiota richness.** As described in Figure 4, we measured peptide levels in supernatants from stable FSA communities cultured in AM, AP, GM, GP, B, and ILC (n=4/medium). Here we report correlations with (A) shared ASV richness and (B) CheSL Shannon evenness. Pearson correlation coefficients and p-values are reported.

**Supplementary Figure S4. Shared richness and mean cross correlation values of FSA communities cultured in AM, AP, B, GM, GP, and ILC across different time points in culture.** As described in Methods, we calculated (A) shared richness and (B) mean cross correlation values for three groupings of time windows during early culture (including the first 3 timepoints A-B), mid (excluding the first three and last three timepoints C-D), and late (excluding the first six timepoints E-F) to identify best time windows for building conserved ensemble networks.

**Supplementary Figure S5. Conserved ensemble networks for FSA and FSB cultured in NS, MM, and IL.** Conserved ensemble networks were built for each media/fecal sample combination from the networks inferred from the window with the highest quality of fit for each replicate. A key describing taxa for each numbered node is provided. Positive interactions are indicated by red lines and negative interactions are indicated by blue lines. (A) Networks for FSA communities grown in NS, MM, or IL as marked below each network. (B) Networks for FSB communities grown in NS and IL as marked below each network. Ensemble networks for FSB grown in MM and 5MM could not be determined due to lack of conserved interactions.

**Supplementary Figure S6. Correlation between carbohydrate diversity and Inverse Simpson diversity for data described by Loss and colleagues.** (A) Fractional data for microbe abundance provided in previously published Supplementary Table 2 was converted to whole numbers and used to calculate Inverse Simpson microbial diversity with mothur v.1.48. Inverse Simpson microbial diversity values were plotted as a function of the carbohydrate diversity measure reported by Loss. Pearson correlation coefficient and p-value are reported. Panels (B) and (C) repeat data from Figure S1 for reference and show correlations between ASV abundance for FSA communities cultured in the indicated media and (B) CheSL Shannon evennes and (C) CheSL Shannon diversity.

**Supplementary File 2: Supplementary Tables S1 and S2**

**Supplementary Table S1: CheSL Richness, CheSL Shannon Evenness, and CheSL Shannon Diversity calculations for media used in this study.**

**Supplementary Table S2: CheSL Richness, CheSL Shannon evenness, and CheSL Shannon diversity calculations for analysis of previously published media described in Supplementary Figure S2.**

**Supplementary File 3: Supplementary Table 3: Table of ASVs generated and analyzed in this study.**

