## Supplementary material for "Diversity in Chemical Subunits and Linkages: A Key Molecular Determinant of Microbial Richness, Microbiota Interactions, and Substrate Utilization": Figure S1 to S6

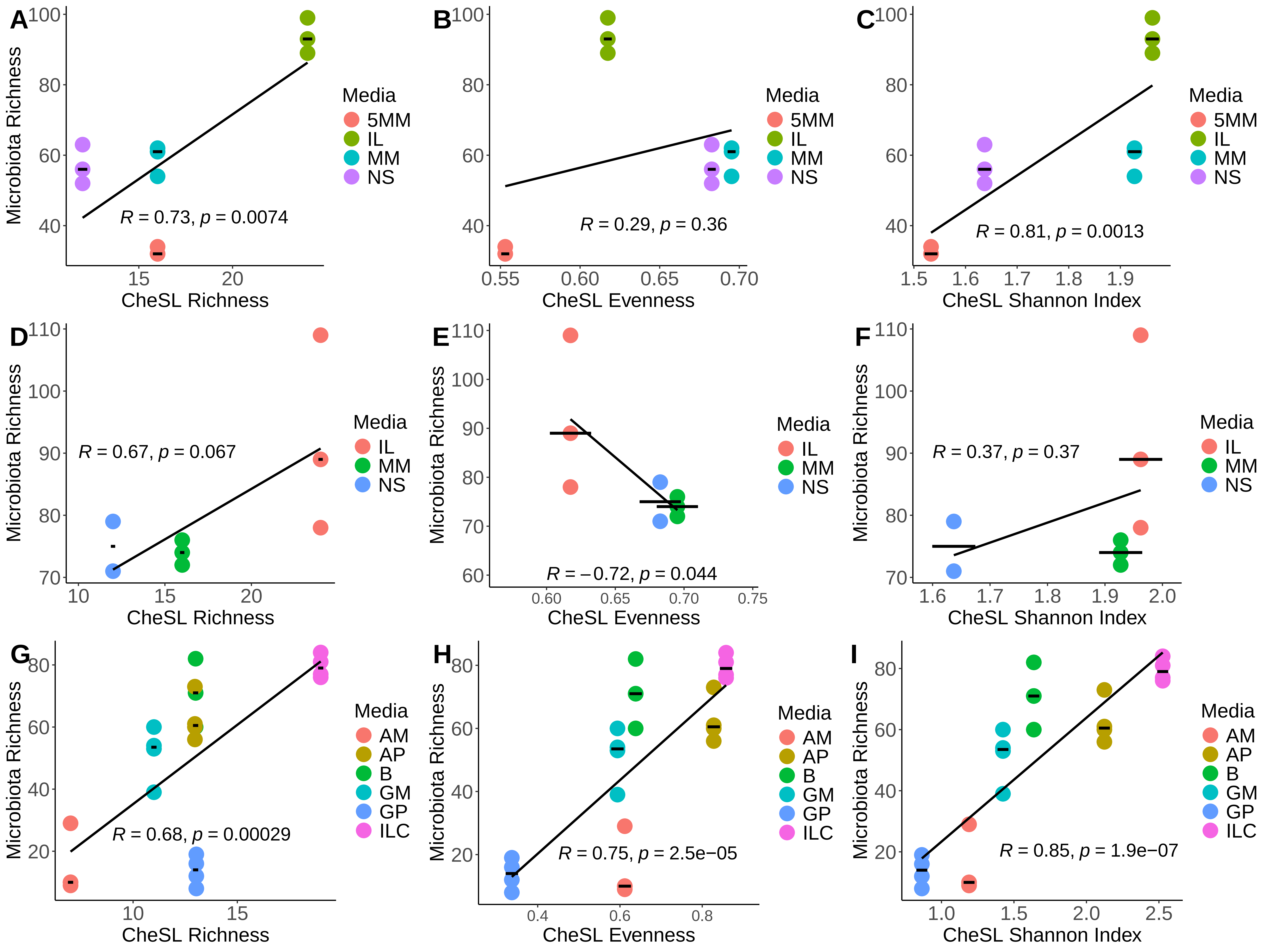

A
Microbiota Richness
CheSL Richness
R = 0.73, p = 0.0074
B
CheSL Evenness
R = 0.29, p = 0.36
C
CheSL Shannon Index
R = 0.81, p = 0.0013
Media
5MM
IL
MM
NS
D
R = 0.67, p = 0.067
E
R = −0.72, p = 0.044
F
R = 0.37, p = 0.37
Media
IL
MM
NS
G
R = 0.68, p = 0.00029
H
R = 0.75, p = 2.5e−05
I
R = 0.85, p = 1.9e−07
Media
AM
AP
B
GM
GP
ILC

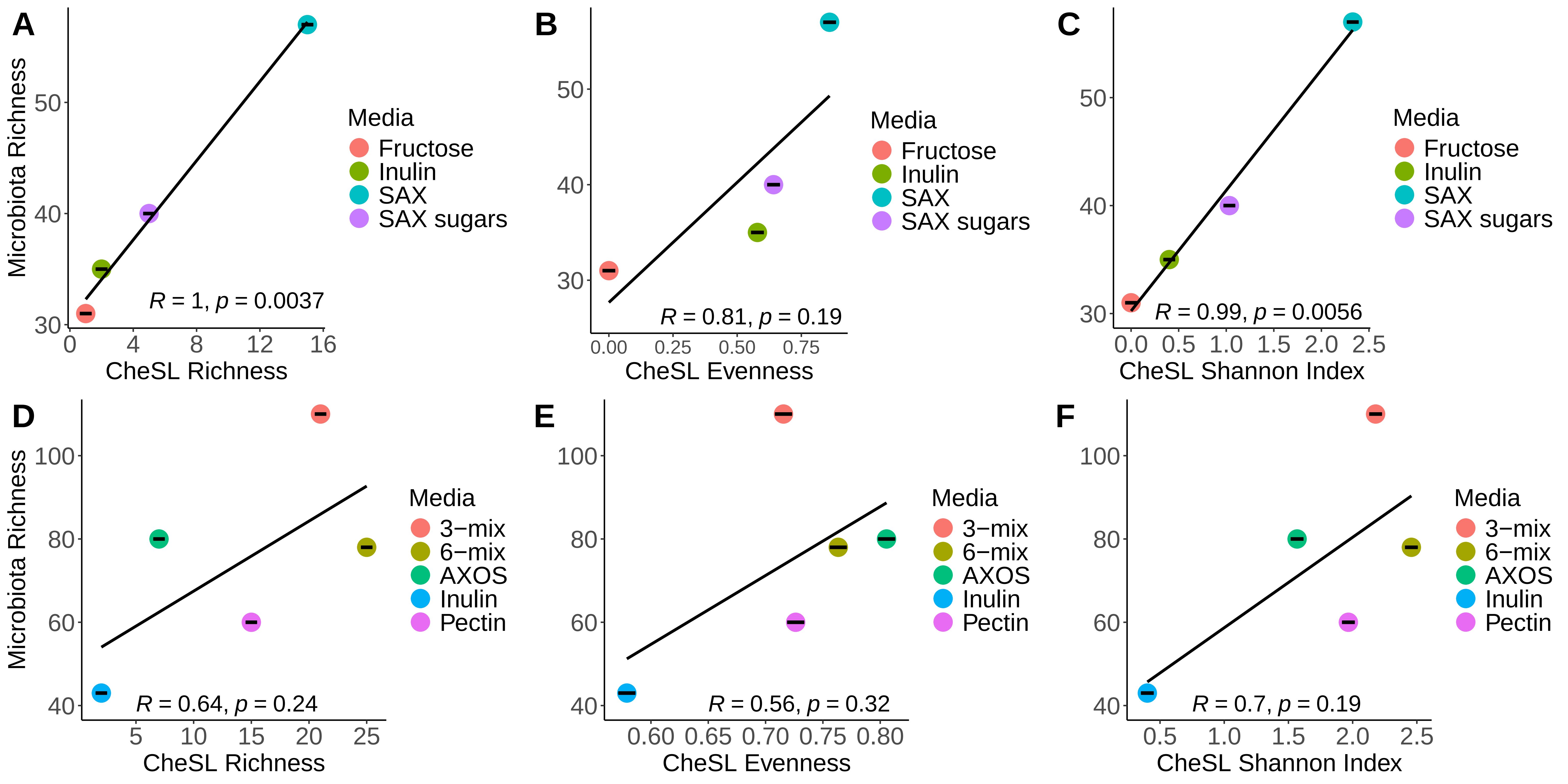

A
Microbiota Richness
CheSL Richness
R = 1, p = 0.0037
B
CheSL Evenness
R = 0.81, p = 0.19
C
CheSL Shannon Index
R = 0.99, p = 0.0056
Media
Fructose
Inulin
SAX
SAX sugars
D
Microbiota Richness
CheSL Richness
R = 0.64, p = 0.24
E
CheSL Evenness
R = 0.56, p = 0.32
F
CheSL Shannon Index
R = 0.7, p = 0.19
Media
3−mix
6−mix
AXOS
Inulin
Pectin

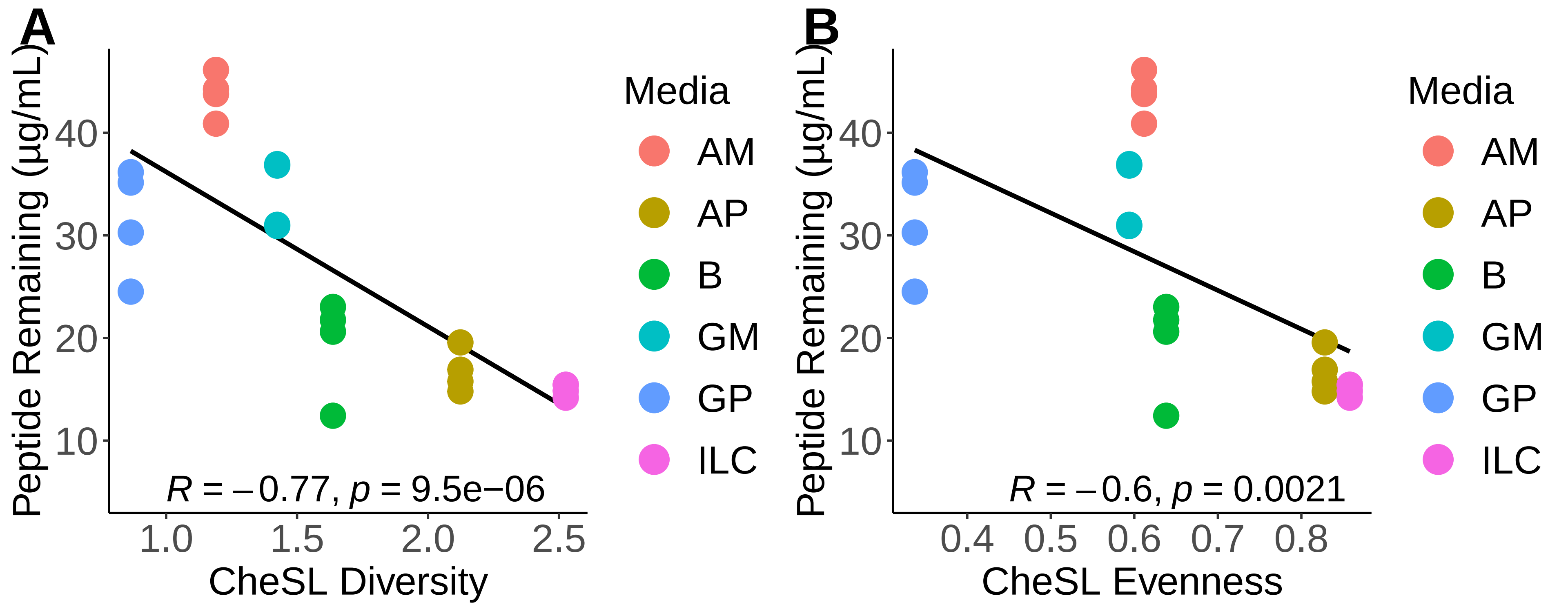

A
B
Peptide Remaining (µg/mL)
40
30
20
10
1.0
1.5
2.0
2.5
CheSL Diversity
R = −0.77, p = 9.5e−06
Media
AM
AP
B
GM
GP
ILC
0.4
0.5
0.6
0.7
0.8
CheSL Evenness
R = −0.6, p = 0.0021

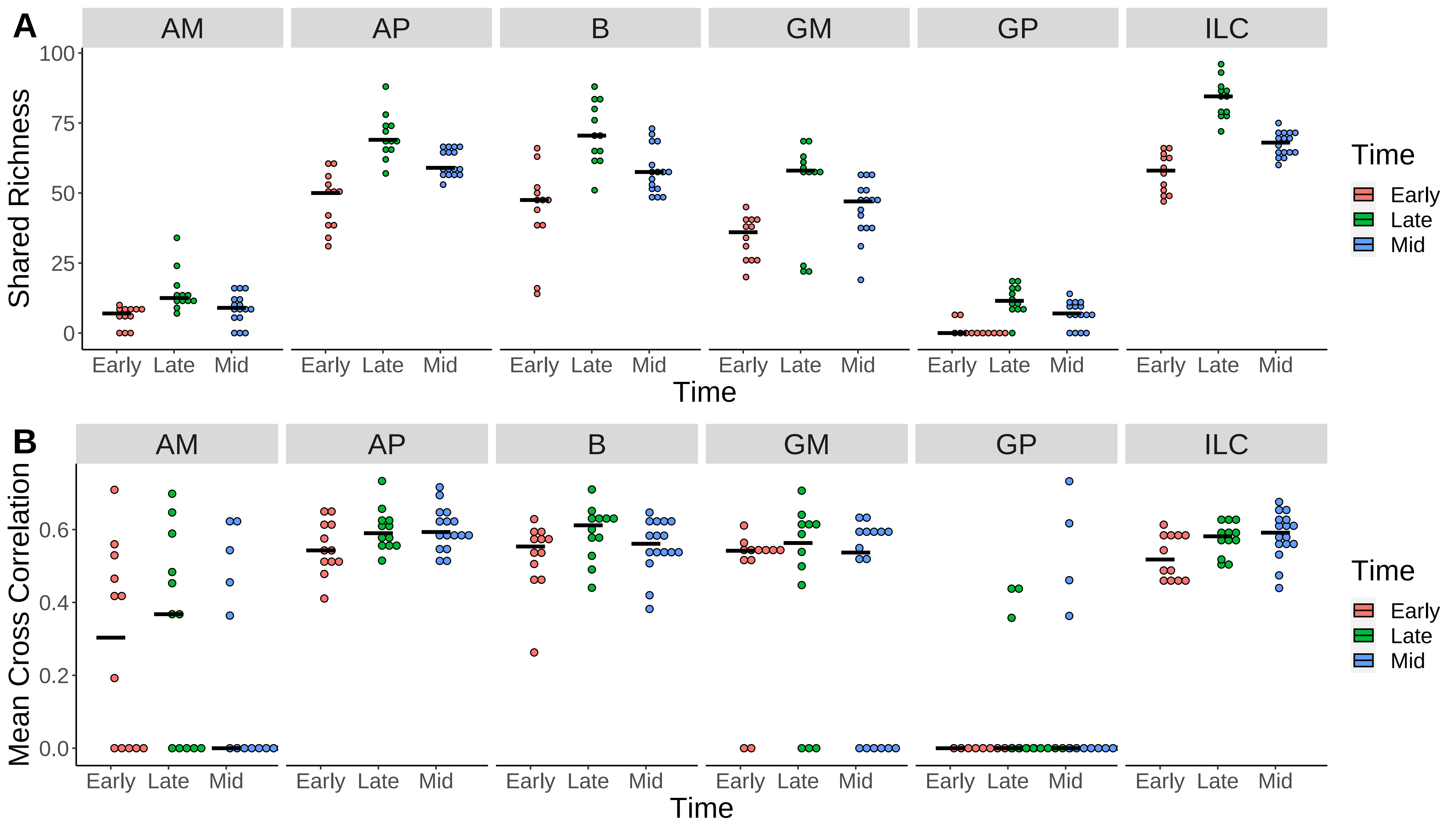
A
B
AM
AP
B
GM
GP
ILC
Shared Richness
Mean Cross Correlation
100
75
50
25
0
0.6
0.4
0.2
0.0
Early
Late
Mid
Time
Time
Early
Late
Mid

**A**

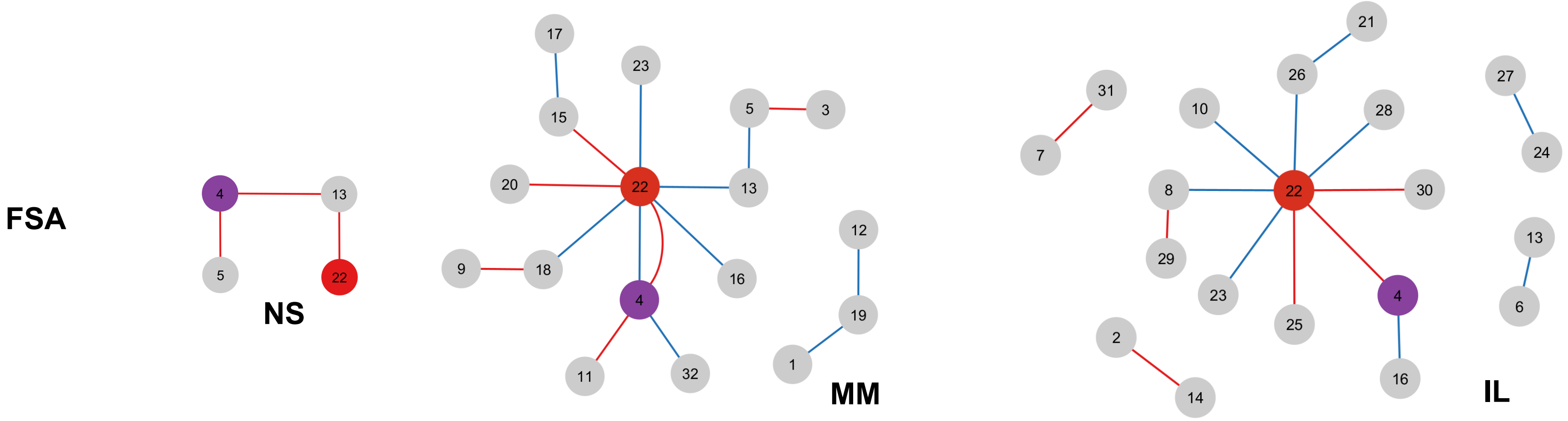


**B**

| Node # | Taxonomy | Node # | Taxonomy | Node # | Taxonomy | Node # | Taxonomy |
|---|---|---|---|---|---|---|---|
| **1** | *Alistipes* | **9** | *Coprococcus* | **17** | *Halomonas* | **25** | *Oscillospira* |
| **2** | *Anaerostipes* | **10** | *Eggerthella* | **18** | *Holdemania* | **26** | *Oscillospiraceae* |
| **3** | *Anaerovoraceae_u* | **11** | *Eisenbergiella* | **19** | *Hungatella* | **27** | *Oscillospirales* |
| **4** | *Bacteroides* | **12** | *Escherichia-Shigella* | **20** | *Incertae_Sedis* | **28** | *Phascolarctobacterium* |
| **5** | *Blautia* | **13** | *Family_XIII_AD3011* | **21** | *Lachnoclostridium* | **29** | *Phocea* |
| **6** | *Christensenellaceae_R-7* | **14** | *Firmicutes* | **22** | *Lachnospiraceae* | **30** | *Ruminococcaceae* |
| **7** | *Clostridia_vadinBB60* | **15** | *Flavonifractor* | **23** | *NK4A214_group* | **31** | *Tissierella* |
| **8** | *Colidextribacter* | **16** | *GCA-900066755* | **24** | *Oscillibacter* | **32** | *UCG-005* |

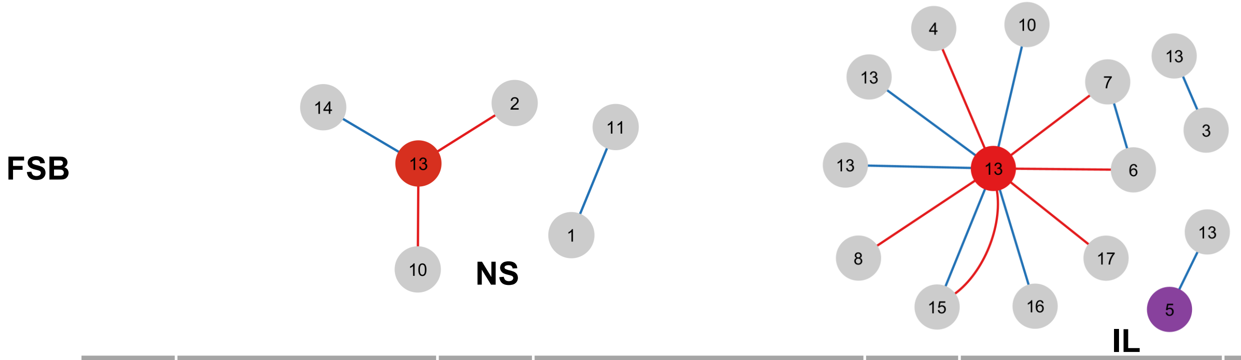


| Node # | Taxonomy | Node # | Taxonomy | Node # | Taxonomy | Node # | Taxonomy |
|---|---|---|---|---|---|---|---|
| 1 | *Acetanaerobacterium* | 6 | *Blautia* | 10 | *Family_XIII_AD3011* | 14 | *Oscillospiraceae* |
| 2 | *Akkermansia* | 7 | *Butyricicoccus* | 11 | *Lachnoclostridium* | 15 | *Ruminococcaceae* |
| 3 | *Alistipes* | 8 | *Christensenellaceae_R-7* | 12 | *Lachnospiraceae* | 16 | *Tissierella* |
| 4 | *Anaerovoracaceae_* | 9 | *Erysipelotrichaceae* | 13 | *Oscillibacter* | 17 | *UBA1819* |
| 5 | *Bacteroides* | | | | | | |

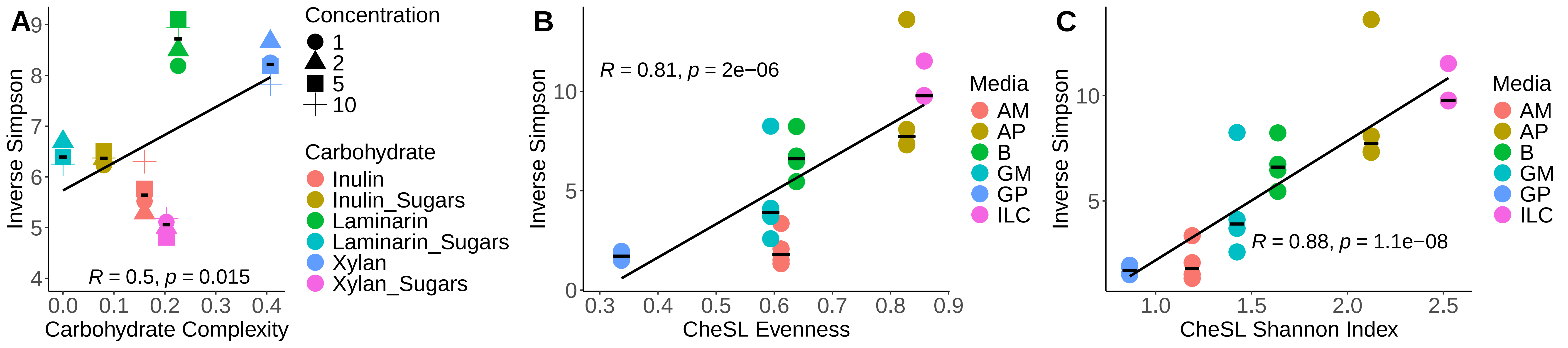
A
B
C
Inverse Simpson
Carbohydrate Complexity
R = 0.5, p = 0.015
Concentration
1
2
5
10
Carbohydrate
Inulin
Inulin_Sugars
Laminarin
Laminarin_Sugars
Xylan
Xylan_Sugars
R = 0.81, p = 2e−06
CheSL Evenness
Media
AM
AP
B
GM
GP
ILC
R = 0.88, p = 1.1e−08
CheSL Shannon Index
